# Identifying Microstructural Changes in Diffusion MRI; How to Circumvent Parameter Degeneracy

**DOI:** 10.1101/2021.09.09.459626

**Authors:** Hossein Rafipoor, Ying-Qiu Zheng, Ludovica Griffanti, Saad Jbabdi, Michiel Cottaar

## Abstract

Biophysical models that attempt to infer real-world quantities from data usually have many free parameters. This over-parameterisation can result in degeneracies in model inversion and render parameter estimation ill-posed. However, in many applications, we are not interested in quantifying the parameters *per se*, but rather in identifying changes in parameters between experimental conditions (e.g. patients vs controls). Here we present a Bayesian framework to make inference on changes in the parameters of biophysical models even when model inversion is degenerate, which we refer to as Bayesian EstimatioN of CHange (BENCH).

We infer the parameter changes in two steps; First, we train models that can estimate the pattern of change in the measurements given any hypothetical direction of change in the parameters using simulations. Next, for any pair of real data sets, we use these pre-trained models to estimate the probability that an observed difference in the data can be explained by each model of change.

BENCH is applicable to any type of data and models and particularly useful for biophysical models with parameter degeneracies, where we can assume the change is sparse. In this paper, we apply the approach in the context of microstructural modelling of diffusion MRI data, where the models are usually over-parameterised and not invertible without injecting strong assumptions.

Using simulations, we show that in the context of the standard model of white matter our approach is able to identify changes in microstructural parameters from conventional multi-shell diffusion MRI data. We also apply our approach to a subset of subjects from the UK-Biobank Imaging to identify the dominant standard model parameter change in areas of white matter hyperintensities under the assumption that the standard model holds in white matter hyperintensities.

## INTRODUCTION

Modelling diffusion MRI (dMRI) data comes in two flavours. Phenomenological models, such as diffusion tensor imaging (DTI) (Basser et al. 1994) and diffusion kurtosis imaging (DKI) (Jensen et al. 2005)) attempt to describe the diffusion signal in a structured mathematical form,while (bio)physical models such as the standard model (Novikov et al. 2019a), NODDI (Zhang et al. 2012), Ball and Rackets (Sotiropoulos et al. 2012) and AxCaliber (Assaf et al. 2008)) attempt to infer properties of the tissue microstructure given the data. This active field of research relies on the inversion of biophysical forward models, but it is also notoriously difficult to overcome model degeneracies (Jelescu et al. 2016). To resolve these degeneracies, the conventional approach is to constrain a subset of the parameters and only make inferences on the remaining parameters (Zhang et al. 2012). However, when the assumptions are not accurate (e.g., in diseased tissue), they will bias the estimated model parameters and cause errors in interpretation. As a result, not only is there a limit to the number of microstructural parameters that can be estimated, but the reliability of the estimated parameters can also be questionable (Jelescu et al. 2016; Reisert et al. 2017; Lampinen et al. 2019).

It is worth mentioning that there are efforts on acquiring complementary information using for example multiple diffusion encoding (Reisert et al. 2019; Coelho et al. 2019; Lampinen et al. 2020), as well as introducing more biophysically informed priors to limit the search space, to provide enough constraints to uniquely estimate the parameters of the standard model. However, here we adopt the standard model of white matter fitted to conventional multi-shell diffusion MRI data as a well-studied degenerate model merely as a toy example to illustrate the concept.

However, in many real-world applications, the model parameters may not be of direct interest. Rather, we are often interested in the “change” in the parameters under different experimental conditions. For example, to study mechanisms underlying a disease one might compare the parameter estimates of biophysical models between patient and control groups. However, the parameter estimation is only tractable when the model of interest is invertible given the data. This limits one to simple biophysical models or requires injection of prior assumptions.

In this work, we show that we can make precise inferences on the change in model parameters even in complex degenerate models. We argue that, using a sparsity assumption on the pattern of change, we can limit the hypothesis space, and so circumvent the degeneracy in the parameter estimation (see Figure 1, also refer to A for more details about directly inferring changes). Our approach proceeds in two steps: First, we use simulated data generated from a forward model to train models that calculate how each parameter affects the measurements. Once these models of change have been trained for all hypothetical patterns of change, we use them to infer the posterior probability of which pattern of change in parameter(s) can best explain the change between real datasets. We call this approach BENCH, which stands for Bayesian EstimatioN of CHange.

**Figure 1.**
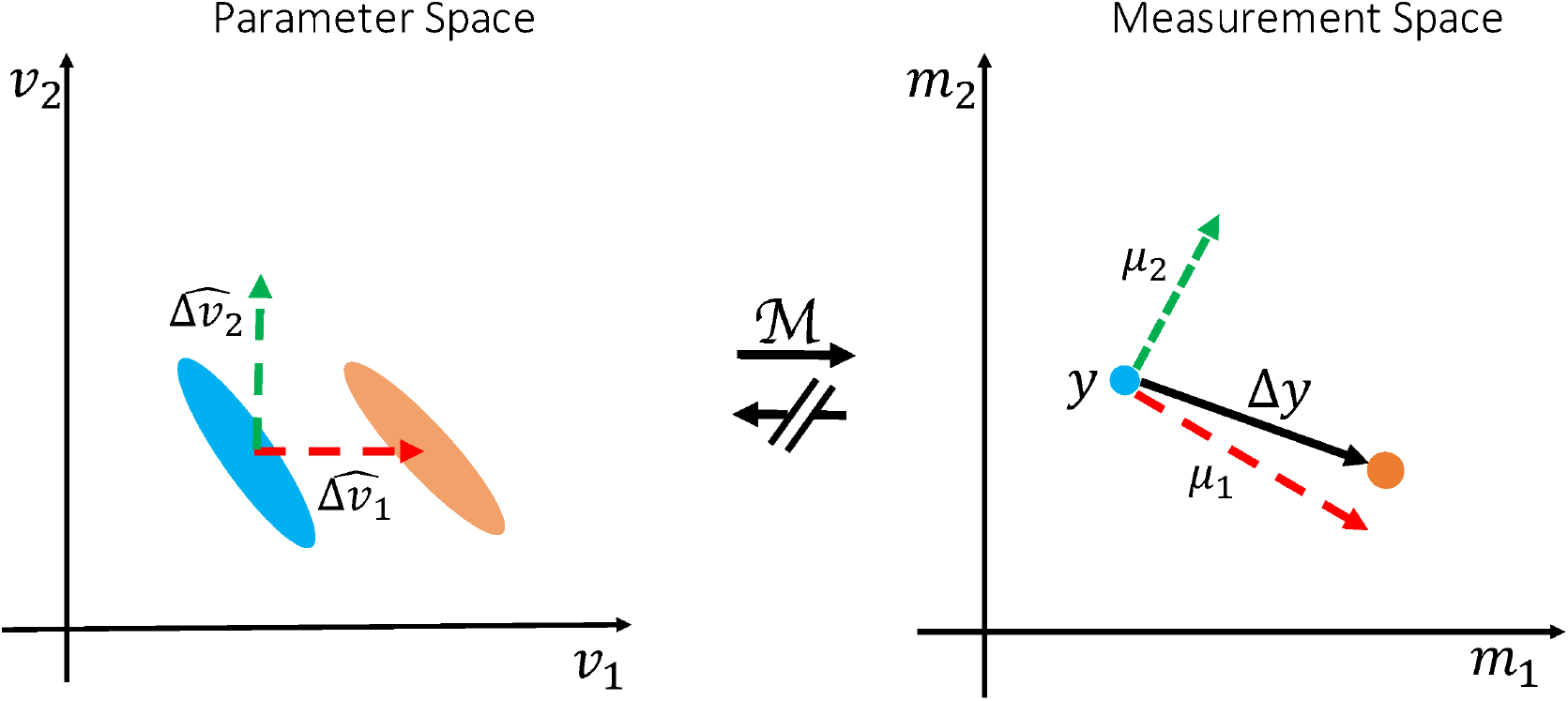
Illustration of the inversion-free inference on change (BENCH). Consider a toy model with two parameters and two measurements ℳ (*ν*_1_, *ν*_2_) = [*m*_1_, *m*_2_]. Each oval in the parameter space (left) corresponds to a single point in the measurement space (right) with the same color; meaning that there is a one to many mapping from measurements to parameters (i.e., the model is degenerate). Despite the degeneracies we are able to estimate which of the parameters best explains the change in the measurements. We do so by comparing the observed change (Δ*y*) with the expected change in the measurements (*µ*_1_, *µ*_2_) as a result of each hypothesised pattern of change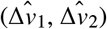.

When confronted with a degenerate biophysical model, BENCH makes a different set of assumptions from the traditional approach of fixing some parameters and identifying any change in the remaining free parameters. When comparing patients and controls, the traditional approach assumes that the prior values for the fixed parameters hold across the region of interest in both groups. Hence, any change of signal across the region of interest between the two groups is assumed to be fully explained by the predetermined set of free parameters. In contrast, by not relying on model inversion, BENCH can work directly with the degenerate biophysical model without fixing any parameters. However, this comes at the price of limiting the change to some predetermined set of possible patterns set by the user (e.g., parameter A could change, or parameter B increases by the same amount as parameter C decreases). While the number of such proposed microstructural changes can be large, each of them has to be sparse (i.e., they have a fewer degrees of freedom than the number of free parameters that could be estimated using the conventional approach). In this work, we will limit ourselves to changes of just one parameter at a time for the sake of simplicity of explanation.

BENCH is applicable to any situation where we are interested in comparing parameters of a generative (bio)physical model across different conditions. Here we apply the framework to diffusion MRI microstructure modelling. As an example use case, we studied microstructural changes in White Matter Hyperintensities (WMH), which are extra bright regions that are commonly seen in T2-weighted images at specific brain regions in elderly people. Despite the abundance and clinical implications of WMHs (Prins and Scheltens 2015; Debette and Markus 2010), the underlying changes in the histopathology and microstructure remain unknown (Wardlaw et al. 2013; Gouw et al. 2011).

The structure of this paper is as follows. In the Theory section, we present the general inference method and how we train the models of change. In the Methods section, we cover the diffusion-specific materials including the computation of summary measurements that are used to represent diffusion data and the microstructural model for diffusion MRI. In the Results section, we first demonstrate the ability of our model in detecting the underlying parameter changes using simulated data. We then apply the method to study microstructural changes in white matter hyperintensities as an example application. In the Discussion section, the potential applications, limitations, and possible future directions of this work are presented.

## THEORY

### Inference on change in parameters

Given a baseline measurement (y), an observed change in the measurement (Δ*y*), and a generative biophysical model (ℳ), we aim to investigate what pattern of change 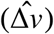 in the model parameters (*ν*) can best explain this observed change in the measurements (Figure 1). A pattern of change is a unit vector in the parameter space, e.g. it can be a change in a single parameter, or any linear combination of the model parameters. For simplicity of the explanations and notation, we only assume a single parameter change in the rest of paper, but all the equations apply to any linear combination of the parameters. If the model is invertible, we may directly estimate Δ*ν* by inverting the model on y and *y* + Δ*y* to get the corresponding parameter estimates and calculate the differences. Alternatively, in BENCH we estimate 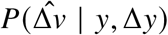, that is the posterior probability for the pattern of change 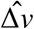 conditioned on the observed baseline (*y*) and change in the data(Δ*y*). Using Bayes’ rule:

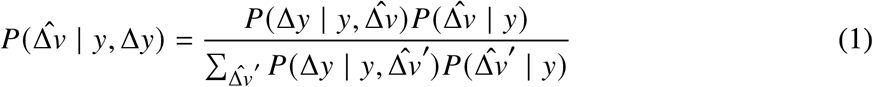

We assume no prior preference between the patterns of change given the baseline measurements(i.e. 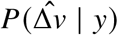 is uniform), so to estimate the posterior probabilities we only need to estimate the likelihood term 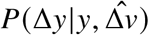. The pattern of change 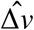 represents the direction but not the amount of the change in the parameters. We therefore marginalize the likelihood with respect to the amount of change (|Δ*ν*|):

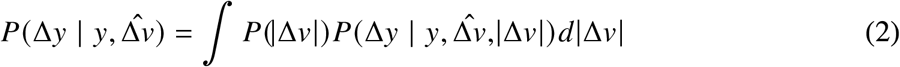

We assume that the prior distribution for the amount of change follows a log-normal pdf with a fixed mean and scale parameter (adjustable hyper parameters). A log-normal PDF is chosen to allow for changes across several order of magnitudes.

The likelihood term inside the integral, 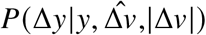, defines how the measurements change as a result of a fully characterised vector of change in the parameters with the given direction 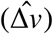 and amount (|Δ*ν*|). To relate this parameter change to a change in data one also needs to know the baseline parameters (*ν*), as

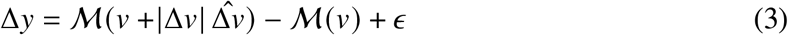

where *ϵ* is the measurement noise. However, for a degenerate biophysical model, we cannot estimate a unique set of baseline parameters *ν* for which to estimate equation 3. While, one could integrate over all possible values of *ν*, this is a very high-dimensional integral, which would be very computationally expensive. Instead, we propose an alternative way to avoid the need of estimating the baseline parameters to estimate the likelihood.

Assuming that |Δ*ν*| is reasonably small, and ℳ is behaving smoothly w.r.t *ν*, using a Taylor expansion we can express Δ*y* as:

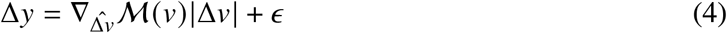

Where 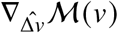 is the gradient of ℳ in the direction of 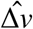 at point *ν*, and *ϵ* is the measurement noise. Given the baseline measurements (*y*), but not the baseline parameters (*ν*), there can be an infinite number of 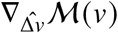 for a degenerate model (Figure 2). To account for all instances of the gradient, we model 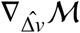 given *y* as a random variable that follows a normal distribution with hyperparameters *µ*(*y*) and Σ(*y*), i.e.

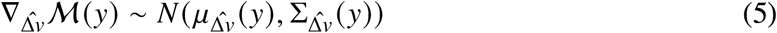

where 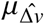 represents the average expected change in the measurements as a result of change in parameters in the direction 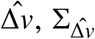 represents the uncertainty around this expectation due to the unknown baseline parameters (Figure 2), and *N* (*m, C*) represents a Gaussian PDF with mean *m* and covariance *C*. This formulation allows us to transfer the uncertainty in the baseline parameters to an uncertainty in the measurement space, which we can model and predict. In the next section we will describe a method for estimating 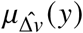 and 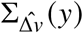 by training regression models using simulated data. Once we compute these hyperparameters, by inserting equation 5 back into equation 4 we can compute the likelihood term inside the integral by

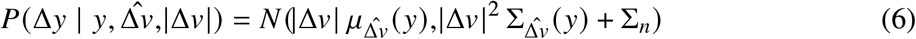

where Σ_*n*_ is the noise covariance matrix.

**Figure 2.**
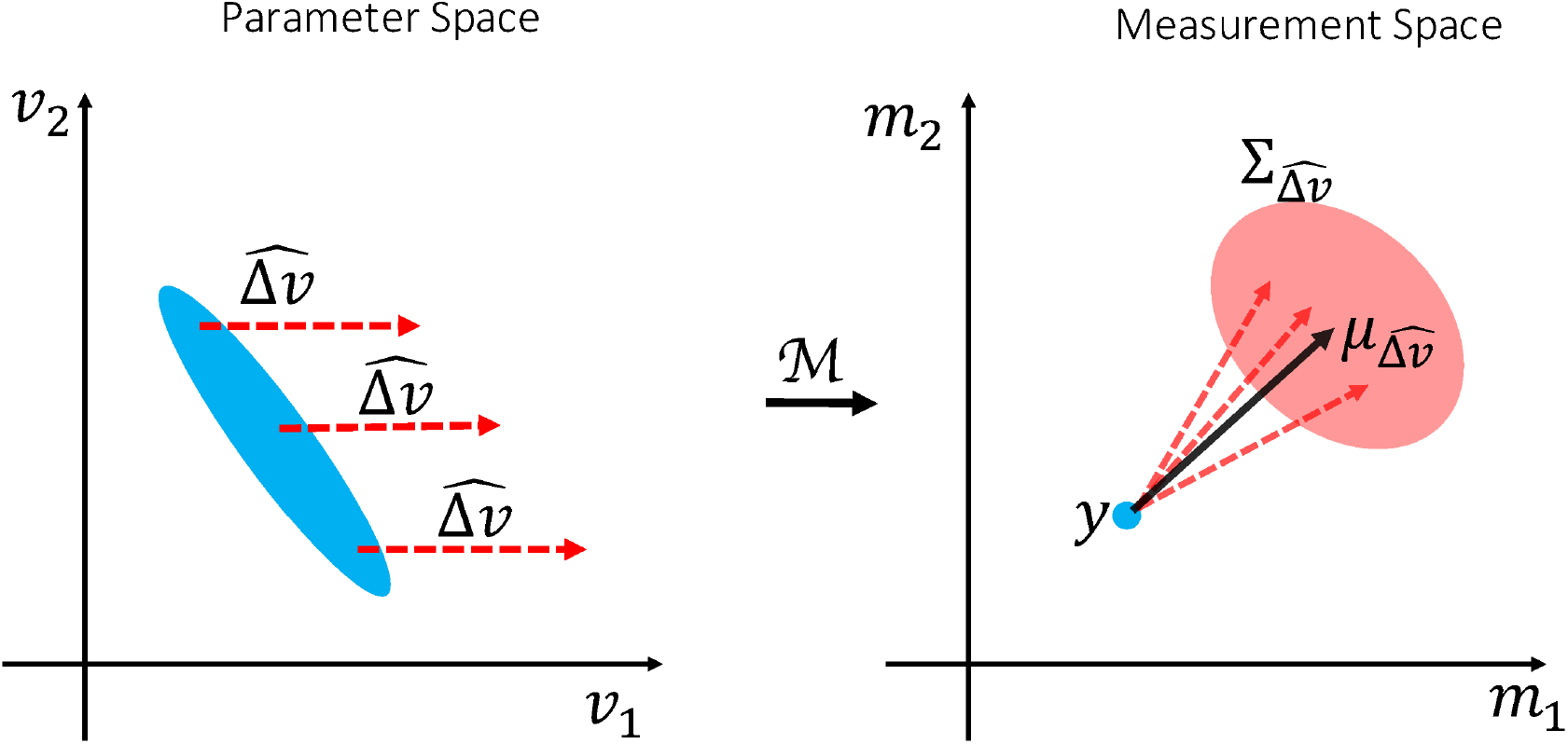
Distribution of gradients. The way measurements change as a result of a particular change in the parameters can only be calculated if we know the baseline parameters. When we are only given the measurements, there are several instances of equally likely gradient directions depending on the underlying baseline parameters. We model all of these gradients given the baseline measurements as a random variable with a presumed distribution. This allows us to transfer the uncertainty due to the inverse model degeneracy into the measurement space. The blue oval in the parameter space (left) represents all the parameter settings that map onto the same blue point in measurement space(right). Each of these parameter settings can produce a different gradient direction in the measurements space. The collection of such gradients of change 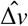for the measurement *y* are modelled as a Gaussian distribution with mean 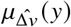 and covariance 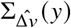.

Finally, by computing the integral over the size of the parameter change in equation 2 numerically, we are able to approximate the likelihood function 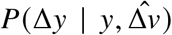 which we can then use in equation (1) yielding the desired posterior distribution on the change in parameters. Moreover, using the approximation of the likelihood function in equation 6 the posterior probability of the amount of change for each direction is proportional to

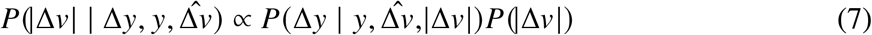

Note that this likelihood function is unnormalized so a high or low values doesn’t necessarily reflect the quality of the change vector in explaining the data. For such measure please refer to appendix B. We can still estimate the most likely amount of change in the parameter given the measurements by finding the|Δ*ν*| that maximizes the above posterior probability (maximum a posteriori estimation). Alternatively, we can estimate the expected value of the amount of change by integrating this posterior probability distribution multiplied by |Δ*ν*| over |Δ*ν*|.

## Training models of change

In this section we describe how to train a regression model to estimate the hyperparameters of the distribution of 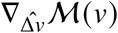, namely the average 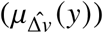 and uncertainty 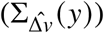 of change in the measurement (*y*) for a parameter change 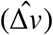.

Given some baseline parameters (*ν*) one can calculate the baseline measurements as *y* = ℳ (*ν*) and approximate the gradient in direction 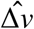 using

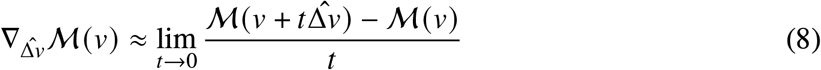

Therefore, by sampling *ν* from the parameter space using a prior distribution, we generate a simulated dataset of pairs 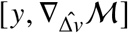 that we use for training regression models.

We use a regression model parameterised by 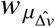 to estimate 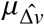 as:

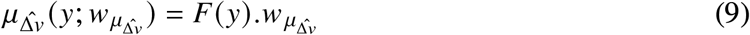

where *F* (*y*) is the design matrix, which depends on arbitrary affine or non-linear transformations of *y*. Note that the subscript 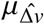 of the weights indicates that each pattern of change in the parameters has its own set of weights.

We also employ a regression model for the uncertainty hyperparameter 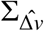 parameterised by 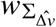. However, 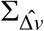 must be positive definite, which would not be guaranteed when directly estimating 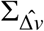 by training an element-wise regression model. To account for the positive definite nature of 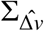, we instead train regression models for elements of the lower triangular matrix of its Cholesky decomposition (*L*). Also, since the diagonal elements of the lower-triangular matrix in Cholesky decomposition must be non-negative, we use their log-transform in the regression model. Hence

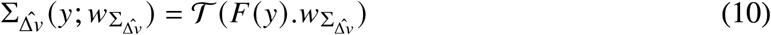

where 𝒯 denotes the transformation of the regressed vector to the full covariance matrix that includes the arrangement of elements, exponentiation of the diagonals, and the matrix multiplication for inverse Cholesky decomposition.

Putting back the above regression models into equation 5 the likelihood of observing pairs of baseline measurements and gradients in terms of the parameters of regression models is:

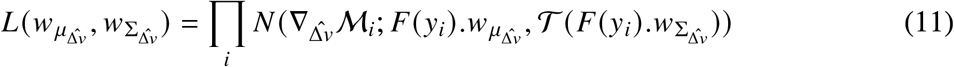

Accordingly, we estimate the optimal weights 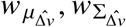by maximizing the above likelihood function for the simulated pairs of 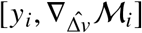 using a combination of the BFGS and Nelder-Mead methods as implemented in SciPy (Virtanen et al. 2020).

This procedure is repeated for each hypothetical pattern of change, yielding two sets of weights for the average and uncertainty of change, which we refer to as a “ change model”. Once we estimated these weights, for any given baseline measurement we use the regression models in equations 9 and 10 to estimate the distribution of derivatives and then the desired probability distributions. Figure 3 shows a schematic overview of the inputs, outputs and steps that are required to train a change model, as well as how to use them to infer the change in parameters.

**Figure 3.**
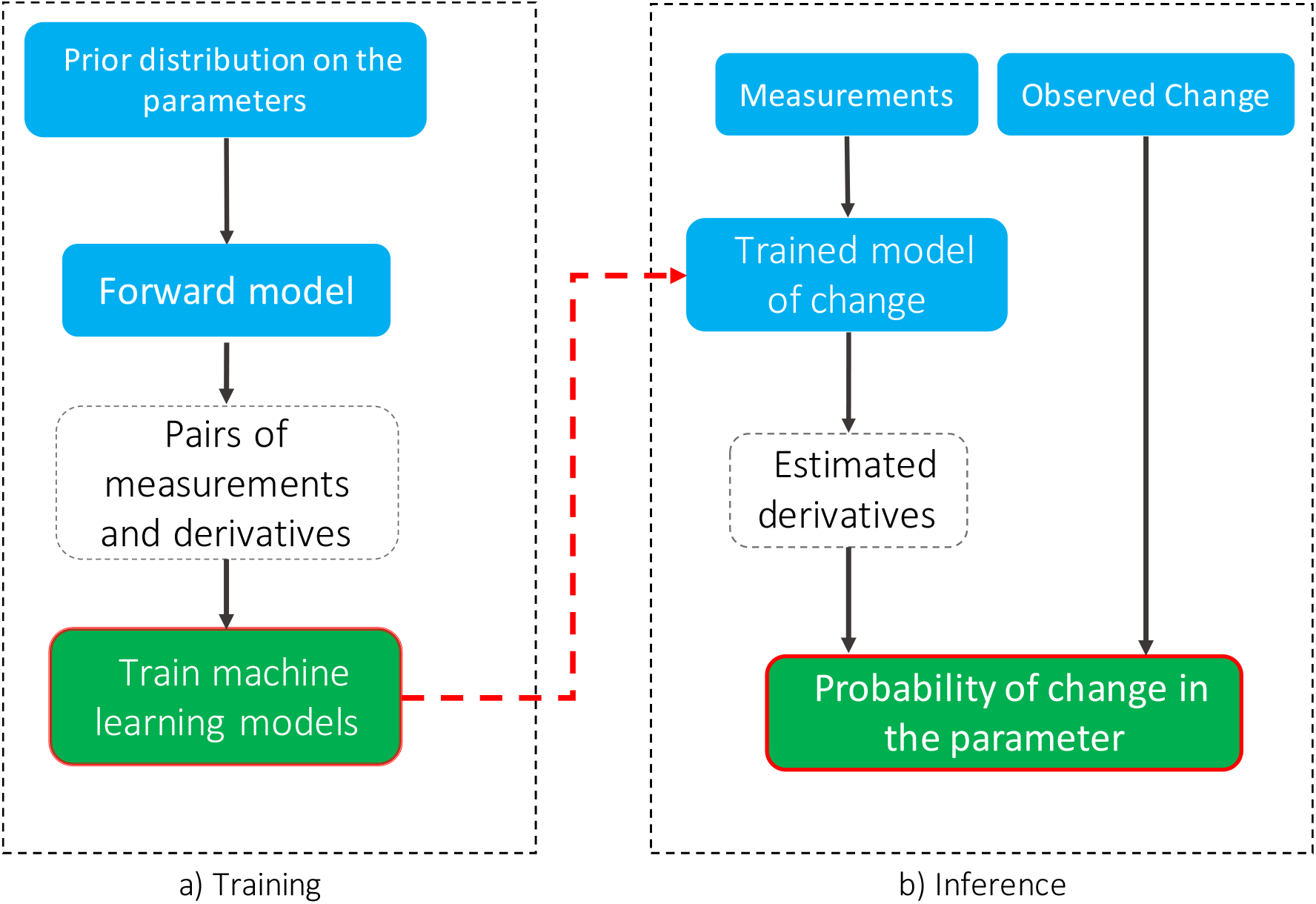
Schematic flowchart for training and inference using change models. The blue, white and green blocks indicate user defined inputs, intermediate variables and outputs respectively. In the training phase for each parameter change, samples that are drawn from the provided prior distribution are passed through the forward model to estimate pairs of measurements and derivatives. Then, regression models are trained to estimate the distribution of derivatives given the measurements using a maximum likelihood estimation. This phase does not require real data and needs to be done only once. In the inference stage using these trained models we estimate the distribution of the derivatives for any given baseline measurements. We then calculate the posterior probability that change in each parameter caused the change in the measurements using the derivative distributions.

In this work, we used a second degree polynomial function of the data for the regression models that estimate the mean change 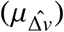 from the baseline measurements. For the uncertainty parameter 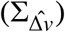a first degree (linear) model is chosen as we expect less variability across samples for this hyperparameter. The weights for the regression models were estimated using a maximum likelihood optimization and a training dataset with 100,000 simulated samples.

### Biophysical model of diffusion

In this section we explain the biophysical model of diffusion that we used to model brain microstructure with diffusion MRI data. The diffusion signal *S* in the brain is conventionally modelled as the sum of signals from multiple compartments. We will here adopt the three-compartment standard model (Novikov et al. 2019a) consisting of an isotropic free water (denoted by the subscript “iso”), an intra-axonal (“in”), and an extra-axonal (“ex”) compartment:

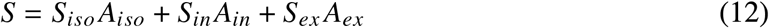

where *S*_*i*_ represents the baseline signal contribution (at *b* = 0), and *A*_*i*_ represents the signal attenuation due to the diffusion weighting in each compartment (Figure 4).

**Figure 4.**
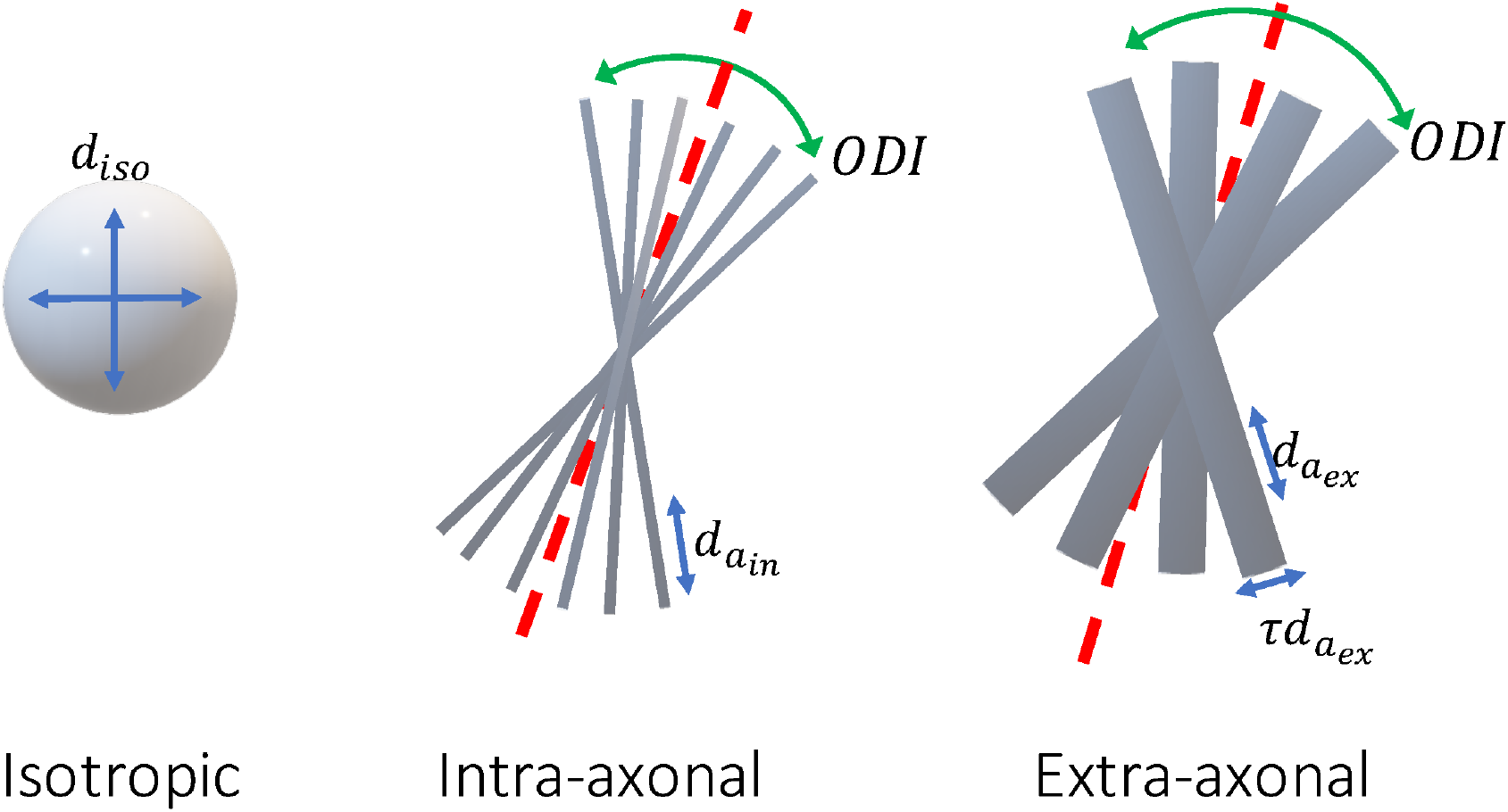
Compartments of the diffusion model. We use a three compartment model that can describe diffusion MRI signals from various brain tissues namely CSF, white matter and gray matter. The isotropic compartment models unrestricted diffusion of water molecules outside of tissue (CSF) with a single free parameter *d*_*iso*_. The intra-axonal compartment models the diffusion of water within axons as several sticks with identical parallel diffusivity parameter *d*_*in,a*_, and zero radial diffusivity, that are dispersed by a Watson distribution with orientation dispersion index *ODI*. The extra-axonal compartment is also a Watson dispersed zeppelin with parallel diffusivity *d*_*ex,a*_ and perpendicular diffusivity *d*_*ex,r*_ = *τd*_*ex,a*_. Including the signal fraction parameters (*s*_*iso*_, *s*_*in*_, *s*_*ex*_) this model has 8 free parameters, which are more than that can be fitted to a conventional dMRI data.

The attenuation for the isotropic compartment is modelled as an exponential decay:

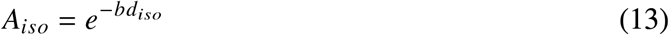

where *d*_*iso*_ is the diffusion coefficient of free water.

The intra-axonal compartment is modelled as a set of dispersed identical sticks with no perpendicular diffusivity. The stick response function for gradient direction *g* and b-value *b* is given by

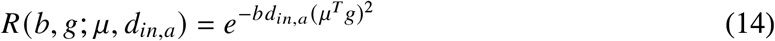

where *d*_*in,a*_ is the diffusion coefficient along the orientation of the stick *µ*.

The fibre Orientation Distribution Function (fODF) is modelled with a Watson distribution, which is defined as

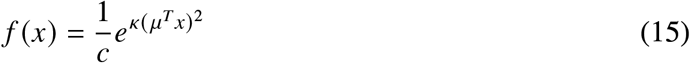

where *µ* is the average orientation, *κ* is the concentration coefficient and *c* is a normalization constant. To assimilate the dispersion coefficient to the notion of variance and limit it to a bounded range, we use the change of variable from *κ* to Orientation Dispersion Index (ODI) as 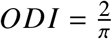 arctan 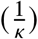. Unlike *κ* which is unbounded, *ODI* is limited to the range (0, 1), where higher *ODI* values correspond to more dispersion. So, the diffusion signal for this compartment is the spherical convolution of the fiber response function with the Watson ODF:

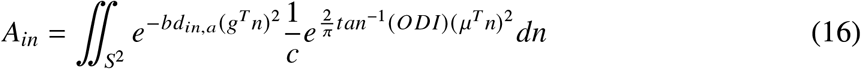

where the integral is over the surface of the unit sphere *S*^2^ representing all possible fibre orientations in 3D.

The extra-axonal compartment is modelled similar to the intra-axonal compartment, with the addition of a non-zero diffusion perpendicular to the fiber orientation. The fiber response function in this case is given by

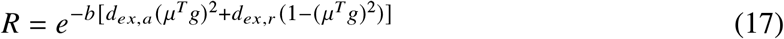

where *d*_*ex,r*_ ≤ *d*_*ex,a*_ are the radial and axial diffusion coefficients. To avoid this dependence between the diffusivity parameters, the parameter *τ* defined as the ratio of perpendicular to parallel diffusivity is used as a substitute to *d*_*ex,r*_. The free parameter τ - subject to *τ* ∈ [0, 1] to maintain the inequality constraint for the diffusivities - can be considered as a measure of tortuosity as it measures the extent to which water diffusion perpendicular to the fibre orientation is hindered with respect to the parallel diffusion. Therefore, the fiber response function for the extra axonal compartment is

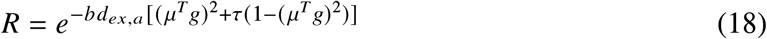

As the compartments share the same geometry, the same fibre orientation distribution is used. Accordingly, the signal attenuation for extra-axonal compartment is given by

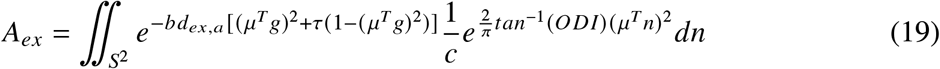

We use the confluent hypergeometric function of the first kind with matrix argument to compute the integrals for both intra and extra axonal compartments similar to (Sotiropoulos et al. 2012).

Table 1 summarises all the free parameters of the described biophysical model along with their valid range.

**TABLE 1.** Microstructural parameters of the diffusion model. All diffusion coefficients are in *µm*^2^/*ms*

| Parameter | Description | Range |
| --- | --- | --- |
| $s_{iso}$ | Signal fraction for isotropic (free water) diffusion compartment | $[0, 1]$ |
| $s_{in}$ | Signal fraction for intra-axonal compartment | $[0, 1]$ |
| $s_{ex}$ | Signal fraction for extra-axonal compartment | $[0, 1]$ |
| $d_{iso}$ | Isotropic (free water) diffusivity coefficient | $[0, \infty]$ |
| $d_{in,a}$ | Parallel diffusivity for the intra-axonal compartment | $[0, \infty]$ |
| $d_{ex,a}$ | Parallel diffusivity for the extra-axonal compartment | $[0, \infty]$ |
| $\tau$ | radial to axial diffusivity ratio for the extra-axonal compartment | $[0, 1]$ |
| $ODI$ | Orientation dispersion index | $[0, 1)$ |

### Summary measurements

Diffusion MRI data are usually measured in multiple shells to capture tissue properties that are sensitive to diffusion of water molecules at various spatial scales. Within each shell, gradients are applied in several directions to measure the geometrical structure of the tissue. However, since we are only interested in the microstructural characteristics, any orientation-related information is irrelevant. We therefore need summary measurements from each shell that are invariant to orientations. We create these summary measurements using real spherical harmonics, which are analogous to the Fourier transform for the spherical domain.

Spherical harmonics are a complete set of orthonormal functions over the surface of a unit sphere. That is to say, any bounded real function that is defined over the unit sphere can be represented by a unique linear combination of these functions with real coefficients. Each real spherical harmonic is denoted by *Y*_*l,m*_ (*θ, ϕ*) where *l* = 0, 1, 2, … is the degree and *m* = −*l*, …, *l* is the order, and *θ* ∈ [0, π], *ϕ* ∈ [−π, π] are the polar and longitudinal angles in standard spherical coordinate system respectively. The diffusion signal at each shell is decomposed as:

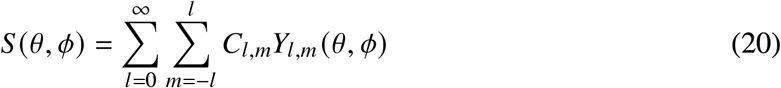

Since the harmonics are a linear basis, one can easily calculate the coefficients for the signal in each shell by inverting the design matrix formed by the harmonics sampled at the gradient directions.

The coefficients are not orientationally invariant. However, the total power in each degree, which is defined as the vector norm of all the corresponding coefficients, is rotationally invariant (Kazhdan et al. 2003; Zucchelli et al. 2020; Novikova et al. 2018). Also, since the diffusion signal is symmetric around the origin and the harmonics of odd degree are odd functions (anti-symmetric w.r.t origin), all odd degrees have zero coefficients.

Consequently, for each shell of diffusion data, we calculate the mean squares of all coefficients for degrees *l* = 0, 2, 4, … as the orientationally-invariant summary measurements.

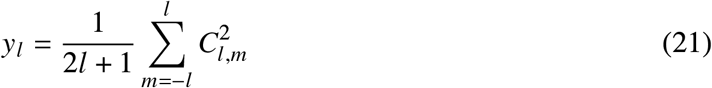

The mean is chosen over the norm to make the scale equal across all degrees. For the case of *l* = 0, we simply use the only coefficient (without the square), so that it represents the mean signal. The higher order summary measurements quantify the signal anisotropy; with greater *l* being more sensitive to sharper changes. We used a logarithm transformation on the power of the coefficients to make the distribution of the measurements for real data closer to a Gaussian and also being more sensitive to smaller changes.

## METHODS

### Simulations

For all the simulations we used the acquisition protocol conducted by the UK Biobank (UKB) (Miller et al. 2016; Alfaro-Almagro et al. 2018) which includes two shells of diffusion 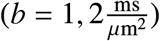 with linear diffusion encoding. Each shell consists of 50 gradient directions distributed uniformly over the surface of the unit sphere, in addition to 5 acquisitions with *b* = 0, yielding a total of 105 measurements.

We used the rotationally invariant summary measurements computed from spherical harmonics for signal representation. The summary measurements for each shell are norms of coefficients at *l* = 0 (absolute value) and *l* = 2 (log mean squared). This produces 5 rotational invariant summary measurements from a diffusion data, namely *b0-mean, b1-mean, b1-l2, b2-mean*, and *b2-l2*.

The described standard model for diffusion is used for both simulated test data and for training models of change. The prior distributions for the parameters are shown in figure 5. We note that these priors are not used for constraining the model parameters but rather they are used to generate training samples for the regression models. The choice of the prior distributions is arbitrary as long as they can reflect all hypothetical parameter combinations that can produce measurements similar to real data.

**Figure 5.**
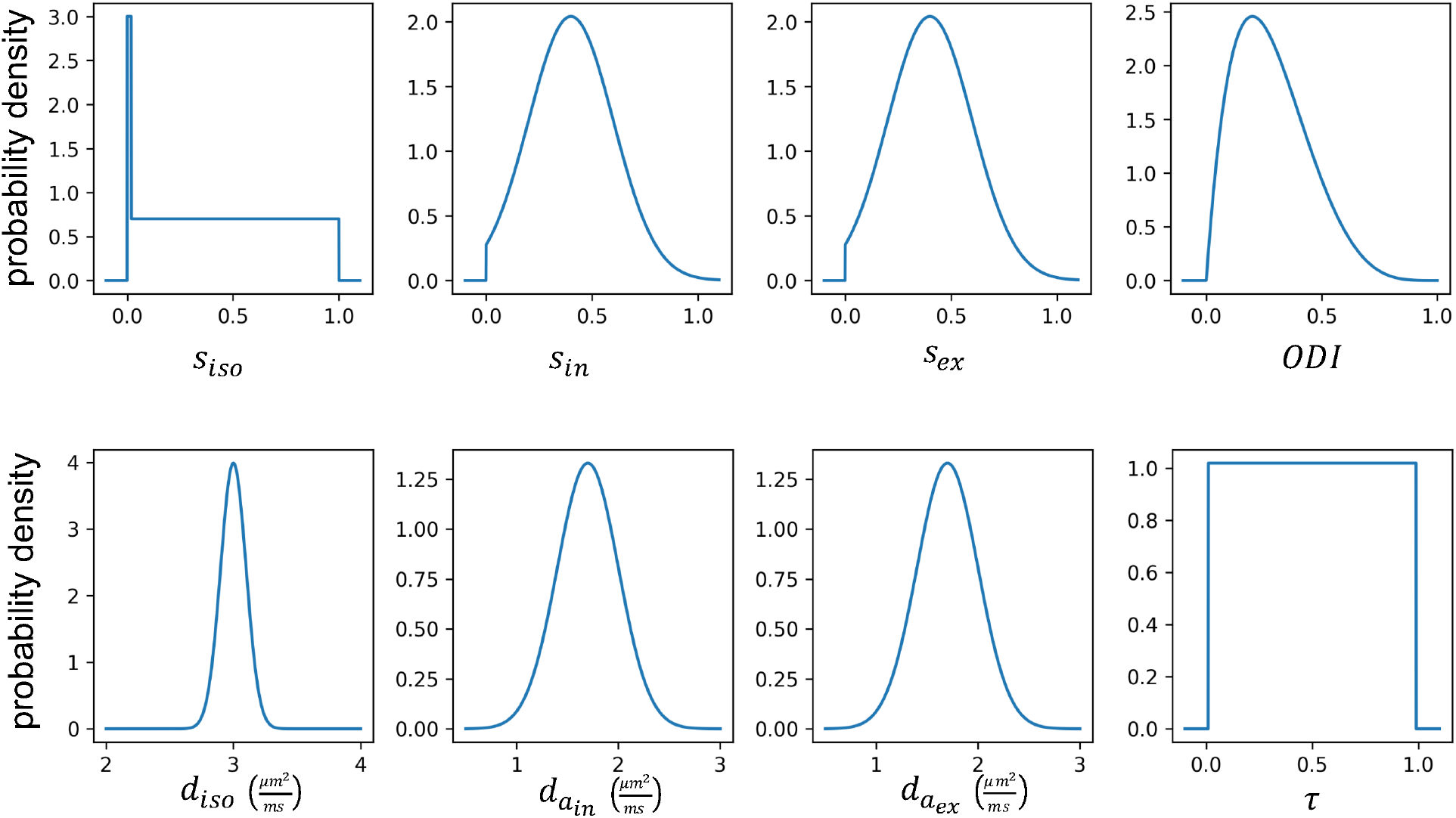
Prior distributions for the parameters of the standard model. These priors are used for generating pairs of measurements and gradients for training the models of change. Also, the same priors are used for simulating test datasets. The priors are chosen such that they contain all probable parameter combinations that can produce measurements similar to real data. The delta function along with uniform distribution in the isotropic signal fraction is used to model pure tissue types as well as partial volume effect. In the training phase, the signal fractions are normalized to sum up to 1. A beta (shape parameters *α* = 2, *β* = 5) distribution is used for *ODI* to impose a nearly uniform distribution for effective fibre dispersion. The prior for isotropic and axial diffusivities are normal distributions with mean 3 and 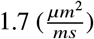 and standard deviation 0.1 and 0.3 respectively; as we expect faster diffusion as well as less variability in the free water component.

The standard model is not invertible given a conventional multishell diffusion data with linear diffusion encoding (Novikov et al. 2019a; Jelescu et al. 2016). Typically, additional constraints are imposed to render the model invertible, e.g. in NODDI (Zhang et al. 2012), the diffusion coefficients are fixed to a prior value as follows:

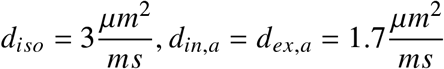

Additionally, the tortuosity parameter *τ* is coupled to the signal fractions:

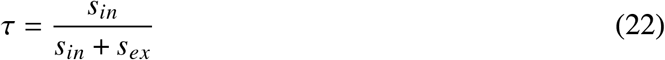

Accordingly, this constrained model has four free parameters: *s*_*iso*_, *s*_*in*_, *s*_*ex*_ and *ODI*.

For both the constrained and unconstrained models, we generated a test dataset containing pairs of simulated diffusion signals, such that in each pair at most one microstructural parameter is different. To generate each pair, we sample a baseline parameter setting from the prior distributions and change one of the parameters by an effect size of 0.1. We also generate pairs of data where no parameter changes and the difference between the two samples is only due to the addition of noise. We then apply the forward model to both parameter settings to produce diffusion MRI signals. Gaussian noise with standard deviation *σ*_*n*_ = 0.01 (SNR=100) is added to all diffusion signals.

The signal fraction parameters are constrained to sum up to 1 for training models of change. Note that whilst this imposes a constraint that the *b0-mean* for the baseline measurement is equal to 1, it does not constrain a *change* in that summary measurement. Accordingly, all the summary measurements (both in the baseline and the change) are normalized by the *b0-mean* of the baseline measurement for any real data. This differs from the parameterization in conventional NODDI, where there is a constraint on the signal fractions to sum up to 1, and add a separate b0 parameter that is directly estimated from b0 signal. Instead, here we assume all the signal fraction parameters can change independently.

For the direct inversion approach, a maximum a posteriori algorithm is employed to estimate the parameters of the constrained model from each diffusion signal separately. Then using a z-test across the parameter estimates in each pair, we calculate a p-value for the change in each parameter (corrected for multiple comparisons across parameters). The parameter with the minimum p-value is identified as the changed parameter. All the cases with minimum *p* > 0.05 are identified as no change.

We also used BENCH for identifying change on the same dataset. To estimate the noise covariance in the summary measurements Σ_*n*_, 100 noisy instances of signals were generated, and the sample covariance of the difference between summary measurements in each pair was estimated. We then estimated the posterior probability of change in each parameter using the trained models of change. The *no change* model has a zero mean and covariance Σ_*n*_ everywhere. The change model with the maximum posterior probability is selected as the predicted change.

### White matter hyperintensities

We investigate the possible microstructural changes in white matter hyperintensities (WMH) using BENCH and model inversion. In this experiment, we used diffusion MRI of 3000 randomly selected subjects from the UK biobank dataset. To account for the variability in overall intensity across subjects, we divided each subject’s diffusion data by the average intensity of the b0 image across the brain’s white and grey matter extracted using FSL FAST (Zhang et al. 2000). We then computed the spherical harmonics-based summary measurements from the diffusion MRI data for each subject and interpolated these measures into the standard MNI space using non-linear transformations estimated by FSL FNIRT (Woolrich et al. 2009; Andersson et al. 2019).

Segmentations of the WMHs were generated from T2 FLAIR images using FSL’s BIANCA (Griffanti et al. 2016) as part of the UK Biobank pipeline (Miller et al. 2016). We computed the average summary measurements for Normally Appearing White Matter (NAWM) that are voxels within the white matter mask not classified as WMH and the WMHs for all voxels that included more than 10 subjects with WMH. For each voxel, subjects were split into two groups according to whether the voxel has been classified as WMH or not. Averaging the summary measures within groups provides us with the baseline measurement (*y*) and the observed change (Δ*y*) related to WMH. The noise covariance (Σ_*n*_) in each voxel was estimated using the within group covariance matrix divided by the number of subjects in the normal appearing white matter group.

## RESULTS

### Summary measurements

A representative axial slice of the normalized summary measurements from a single subject are shown in Figure 6. The *“mean”* summary measures represent the normalised average signal. The *l*2 measures quantify the anisotropy in each voxel (similar to Fractional Anisotropy maps in DTI).

**Figure 6.**
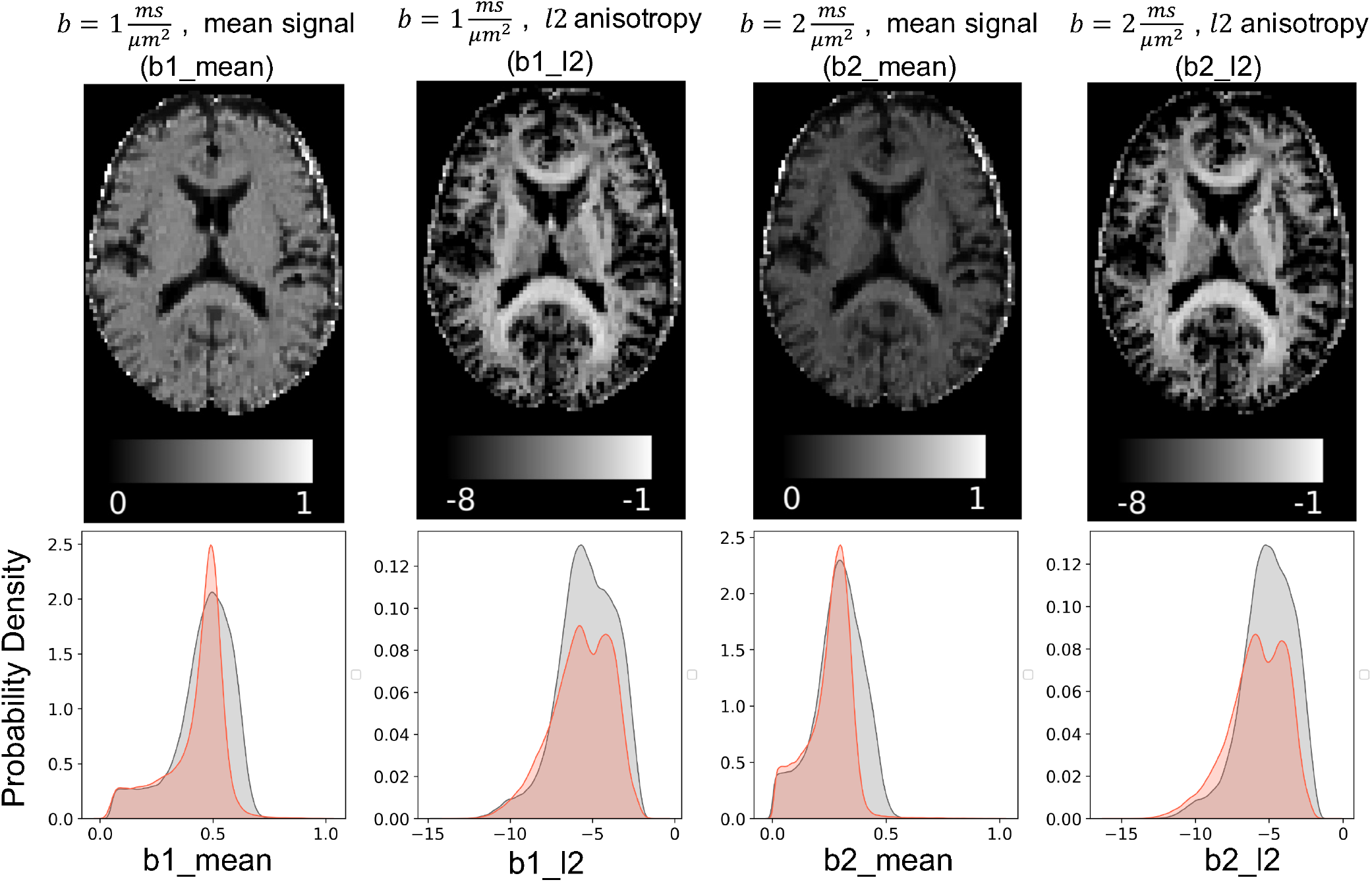
Maps of the summary measurements for a sample subject in the UK biobank dataset (top) and their histogram (bottom). The mean summary measurements is reflecting the average (across directions) diffusivity in each shell. The *l*2 summary measurements estimate the anisotropy, which is similar to the fractional anisotropy (FA), but computed with a linear transformation of the signal. Histograms show the distribution of these measurements across the brain; as well as the distribution of simulated data using the standard model and provided prior distributions. This shows that the simulations capture the full range of the summary measures from real data.

The bottom panels of Figure 6 show histograms of the summary measurements across the brain for the same subject, as well as distributions of simulated data based on prior distributions over the model parameters. The distribution for the generated samples fully covers the range of the data and follows a very similar density distribution. This verifies that the prior distributions are wide enough to capture the full range of real data.

Figure 7 shows estimated derivatives of the summary measurements at baseline data representative of putative voxels in the white matter and grey matter. The error bars show estimated standard deviations of the derivatives (the square root of diagonals of the estimated covariance matrix). This variance is reflecting the uncertainty in the underlying parameters that can generate these measurements, as well as residuals of the regression model for the mean.

**Figure 7.**
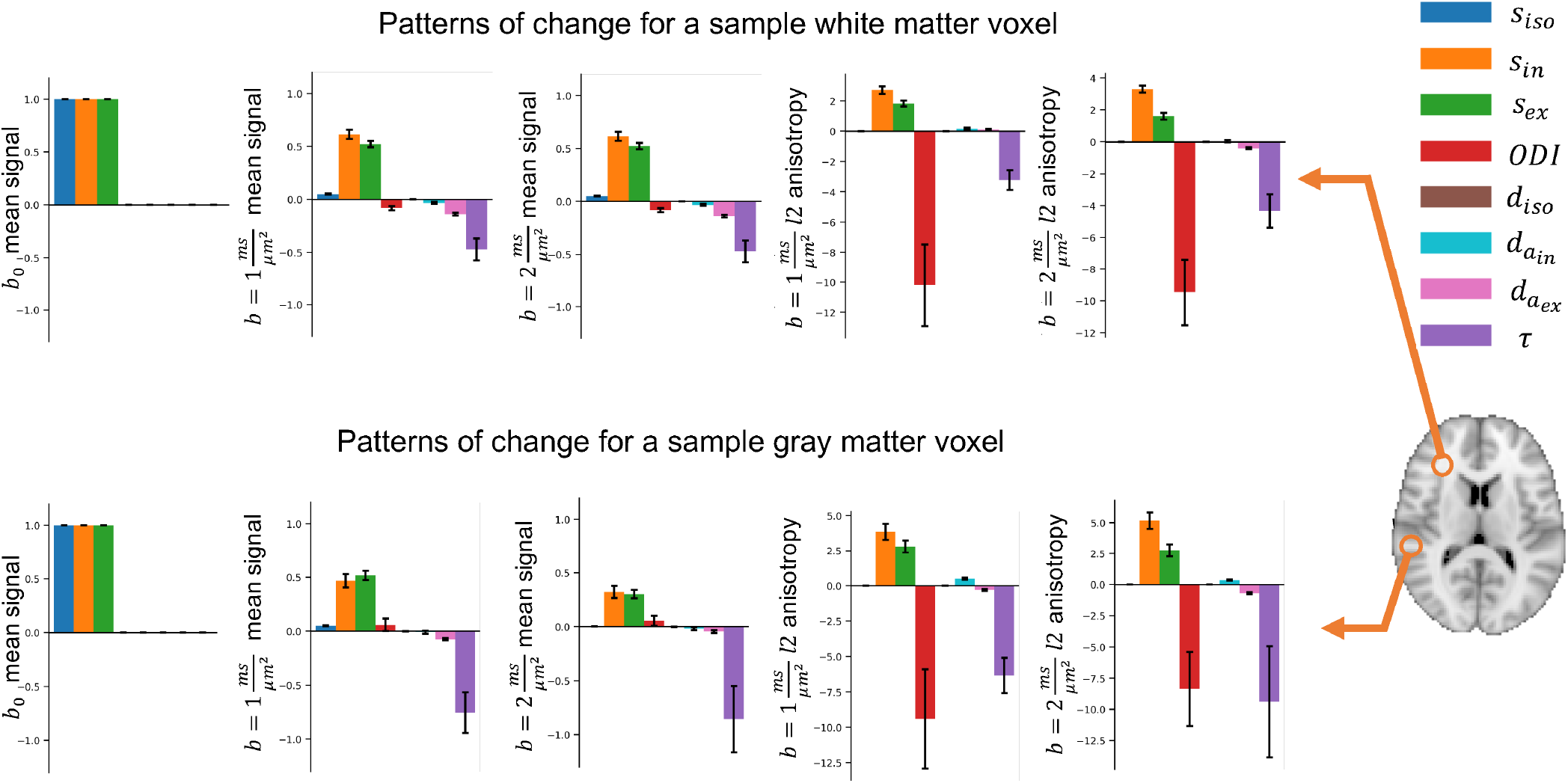
The estimated amount of change in the summary measurements as a result of a unit change in each parameter 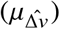for a sample white matter and grey matter voxel. The error bars show the estimated standard deviation of change. Colors correspond to parameters and columns indicate summary measurements. Due to differences in the baseline, each voxel can have a different change vector for the same parameter change. This added degree of freedom can model the variability of parameters (e.g. diffusivities) across the brain, which is not considered in constrained models; e.g. NODDI.

### Validation

We first employed simulated data to evaluate the performance of the proposed approach in inferring microstructural changes from diffusion MRI data. The details of experiment parameters are provided in the methods section.

#### Comparison with model inversion

Figure 8a shows the confusion matrix using model inversion (left), and our inversion-free approach (right) for an invertible model with only 4 free parameters. Each element of these matrices represents the percentage of times a change in the parameter represented at the corresponding column is identified as a change in the corresponding row. Both approaches were able to detect the true parameter change in most of the cases.

**Figure 8.**
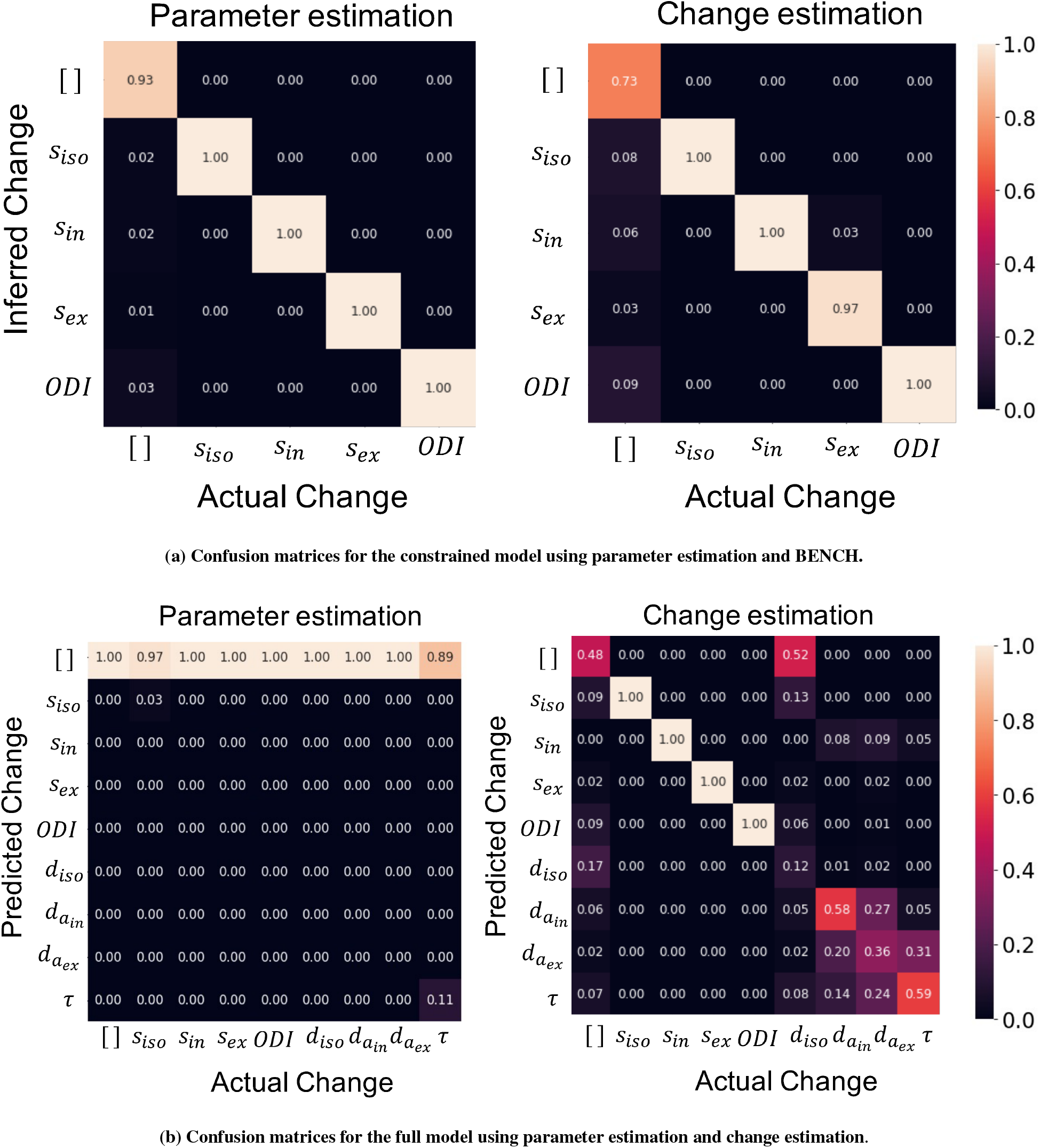
a) The numbers indicate the percentage of time a change in the corresponding column is identified as a change in the corresponding row. The diagonal elements show the accuracy in identifying true change. a) Both of the approaches performed near to ideal in detecting the true change in the case of constrained model. The change estimation has more false positives, but unlike the inversion approach, we did not explicitly define a false positive rate threshold. b) Given diffusion data at few shells, the full model is not invertible, i.e. the parameter estimates have a high variance. Therefore, almost no significant change is detected using parameter estimates. On the other hand, the change estimation approach can still identify changes in all the parameters of the restricted model. Although there remains confusion between a subset of the parameters since these have similar effects on the diffusion signal.

For the standard model with all 8 free parameters, Figure 8b shows the confusion matrices using the direct model inversion (left) and change estimation (right). Since the uncertainties of the parameter estimates are very large due to the model degeneracies, almost all of the changes are confused with *no change* when using direct inversion. However, the inversion-free approach is able to identify changes in *s*_*iso*_, *s*_*in*_, *s*_*ex*_ and *ODI*. Although, there is confusion between the remaining parameters compared to the restricted model, here we do not make any strong assumptions on the value of those parameters. Also, most of the confusions for these parameters are between them, meaning that we are able to distinguish a change in those parameters (e.g. the diffusivity parameters) from others. Change in isotropic diffusivity is mostly confused with the *no change* model. This is due to the *b*-values in the UKB protocol which are too high for this parameter; a change in this parameter has minimal effect on the signal.

#### Sensitivity to change in each parameter

To evaluate the sensitivity of the approach to the amount of change in each parameter, we generated test datasets with variable effect sizes starting from 0 to 0.1 with step sizes of 0.01. Figure 9a shows the average posterior probability of change in each parameter versus the effect size. In all types of change, at very small effect sizes (< 0.01) the change is confused with no change, but as the effect size increases the probability of identifying the true change (red curves) increases. Changes in all signal fraction parameters and in the fibre dispersion are identified with high accuracy even at very small effect sizes. However, changes in diffusivity parameters are confused with each other (but not with signal fraction parameters) even at larger effect sizes. It is worth mentioning that effect size and SNR are two important factors (both unknown in real data) that affect the performance of detection in a similar way. So, when SNR is lower (resp. higher) the approach can be more (resp. less) sensitive to the change. Here we show the results for SNR=100.

**Figure 9.**
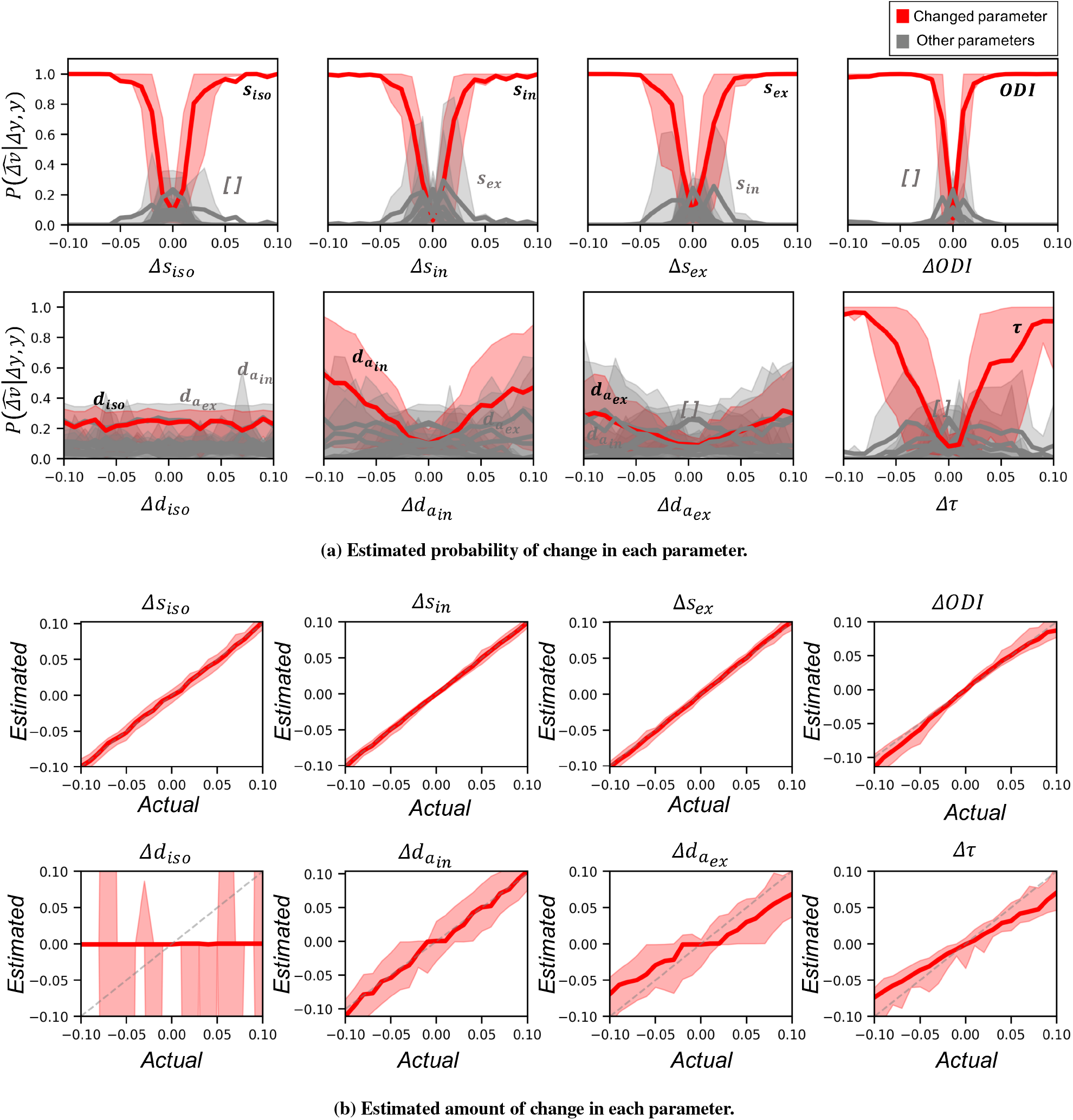
**a)**Each plot shows the estimated probabilities when the corresponding parameter on the *x* − *axis* has changed between two datasets. Red curves show the average posterior probability of change in the actually changed parameter versus the amount of change. The gray curves show the probability for other parameters. Shaded areas show the 10 to 90 percentile range. Larger absolute amount of change results in higher posterior probability for the true parameter change. Change in the signal fraction parameters and *ODI* is distinguishable for effect sizes as small as 0.05. However, changes in diffusivity parameters even at very large effect sizes is cluttered with other parameters. **b)** Each plot shows the maximum a posteriori estimation of the amount of change vs the actual change in the parameter. The shaded areas show the 10 to 90 interval. The estimated change in the signal fractions follow the identity line (dashed gray line). The estimated change in *d*_*iso*_ is mostly around zero with a high variance as the posterior distribution is very flat and symmetric around zero. The change in *d*_*ex,a*_*τ* and *ODI* is systematically biased at higher effect sizes.

#### Estimating the amount of change

So farwe have only examined the posterior probabilities relating to the identity of the parameters that can best explain a change. However our framework also allows us to estimate the posterior probability on the *amount* of change for each parameter 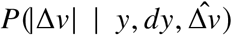 (eq.7). Figure 9b shows the estimated (maximum a posteriori estimation) versus actual change in each parameter for different effect sizes.

### White matter hyperintensities

#### Model Inversion

We inverted the NODDI model using non-linear fit implemented in DMIPY (Fick et al. 2019) in all subjects and ran a voxel wise glm to estimate the differences between white matter hyperintensities and normally appearing white matter (NAWM). Unlike in BENCH, NODDI requires fixing the diffusivity parameters. Usually, they are fixed to 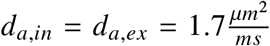 (and 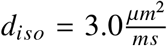). However, it has been recently suggested that the axial diffusivity should be higher based on several studies attempting to directly measure their value (Howard et al. 2020; Kunz et al. 2018). We have therefore run the same analysis also with 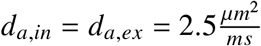.

The z-maps for the contrast of WMH vs the baseline for all the parameters are shown in Figure 10. The strongest changes are seen in *f*_*intra*_ and it is consistent in both high and low diffusivity regimes. The direct inversion also suggests changes in the other two parameters (*f*_*iso*_ and *ODI*). However, interestingly, changing the pre-specified diffusivities in NODDI alters the story for *f*_*iso*_ and *ODI* which go in opposite directions(see scatter plots in 10 many points(voxels) lie in the 2nd or 4th quarter). These results demonstrate that the choice of fixed parameter values can affect the inferred change in other parameters.

**Figure 10.**
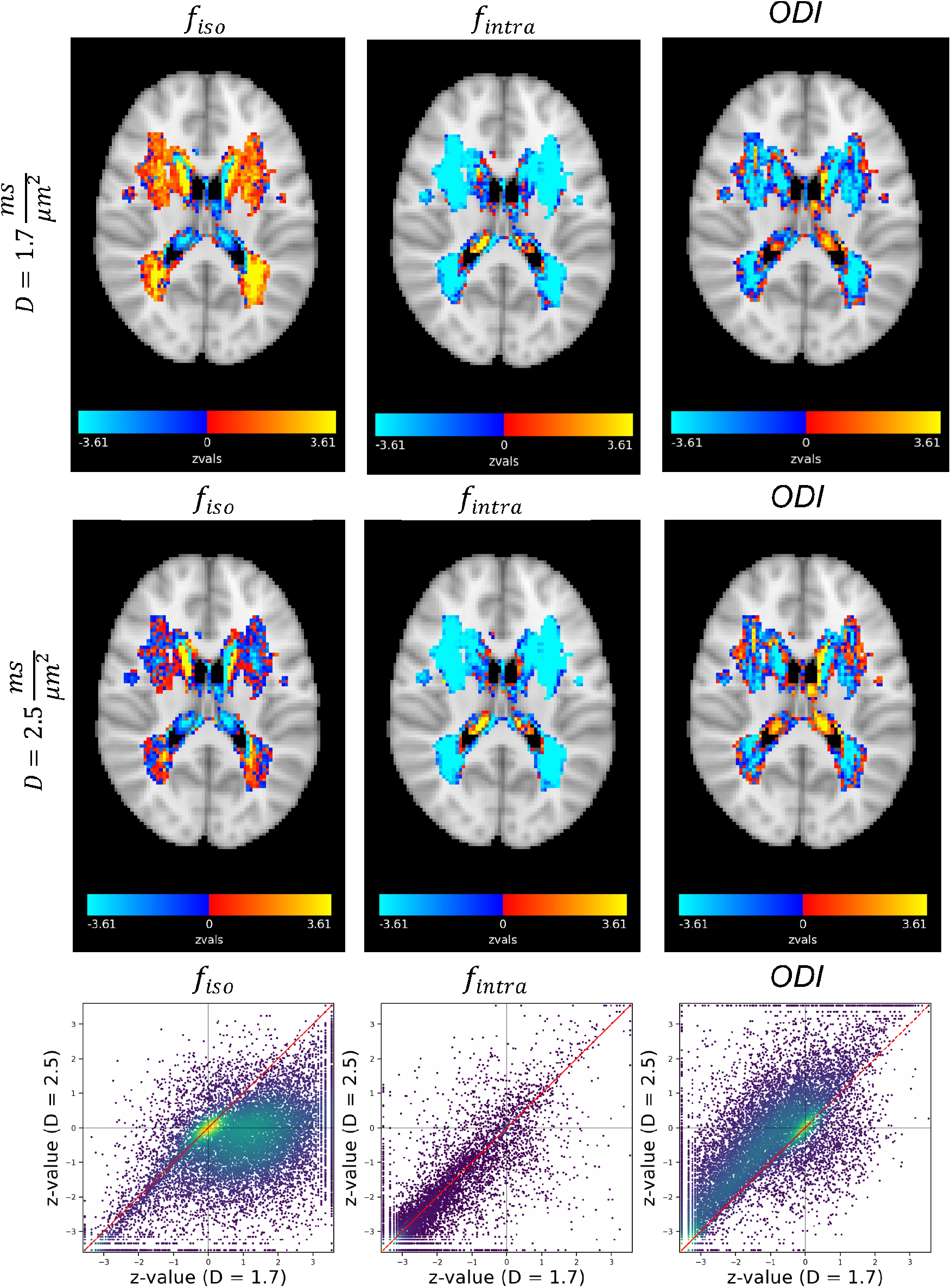
NODDI parameter estimates. **Top)** z-maps for the difference between WMH and normally appearing white matter with the assumption 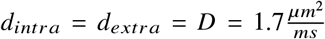. **Middle)** The same maps with the assumption 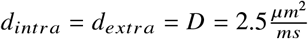. **Bottom)** Scatter plot of the z-values for the two cases. The results show the assumed fixed value for the diffusivity significantly affects the estimated change between WMH and normal tissue for *f*_*iso*_ and *ODI*. However, the observed decrease in *f*_*intra*_ is fairly robust to the difference in diffusivities, and this is inline with the results from BENCH.

#### BENCH

We used the trained models of change on the parameters of the full (standard) model to infer changes in WMH. Figure 11a shows the observed change in the summary measurements (normalized by b0 mean of the baseline) in white matter hyperintensities (dashed line) as well as predictions from each model of change(colored bars) for average data from a small patch of white matter. For each parameter, the best amount of change given the baseline, the observed change and the noise covariance is estimated using equation 3. In other words, the bars indicate the closest change in the measurements that can be produced when only that parameter has changed.

**Figure 11.**
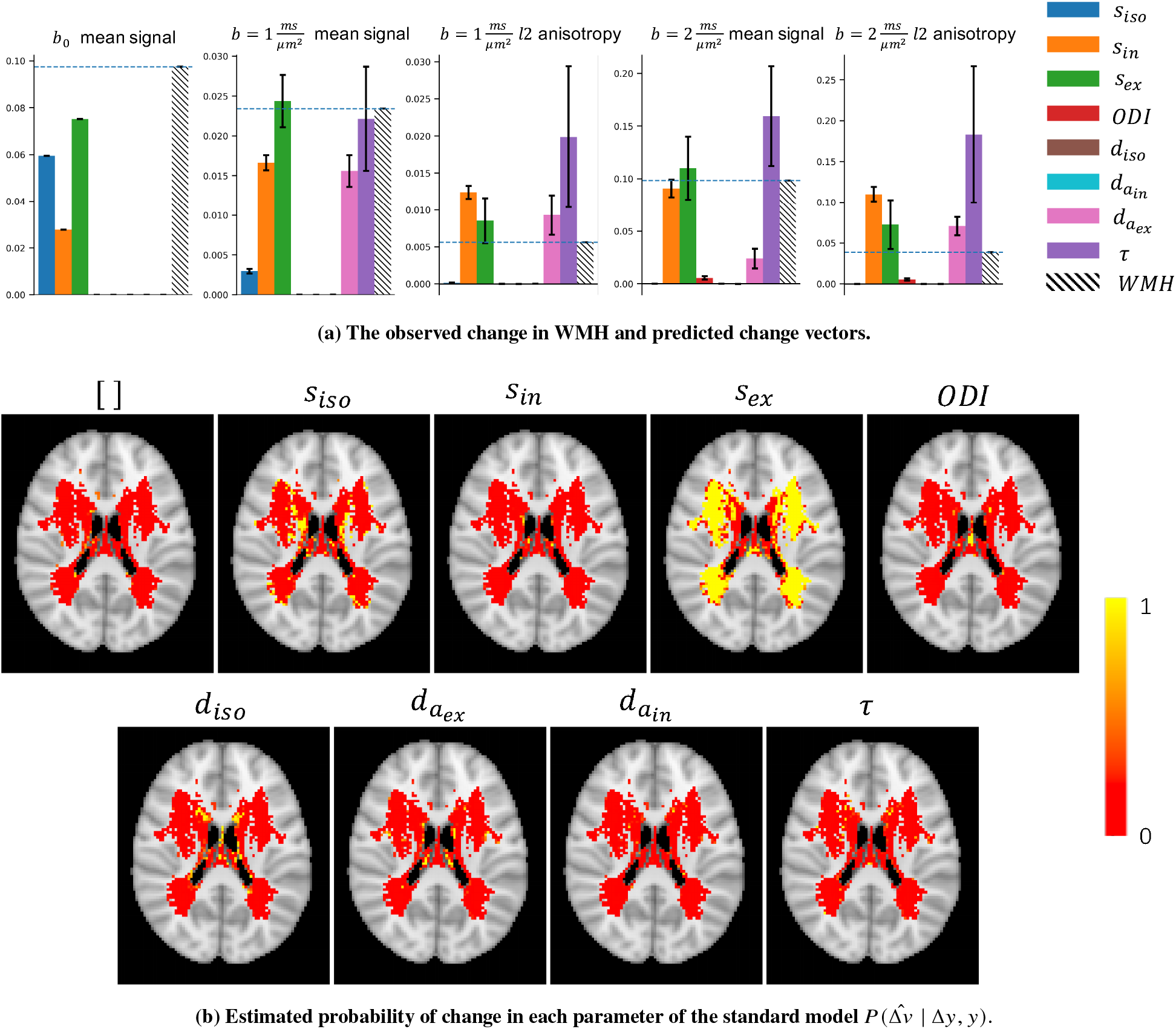
**a)** Each panel shows the estimated amount of change in the measurements if only the corresponding parameter changes, along with the actual observed change in hyperintensities for a patch of voxels in white matter. Each bar is scaled with the best estimated amount of change for that parameter. The observed change in WMH is an increase in the *mean-b0* and, to a lesser extent, and increase in *mean-b1*, and a positive change in the l2 measurements. This is best aligned with the pattern of change that an increase in *s*_*ex*_ can produce. **b)** Each map shows the estimated probability that change in the corresponding parameter can explain the observed change in the summary measurements between WMH and NAWM at a single axial slice of the brain. The no change model represents the null hypothesis that the change is better explained by noise rather than a change in any one of these parameters. In the majority of the voxels, the change model for *s*_*ex*_ has a probability around 1 (yellow) and the remaining parameters are nearly zero(red). This means that a change in *s*_*ex*_ is more likely to explain the observed change than any other single parameter change.

This plot suggest that the observed change in WMH is an increase in the *b0_mean* and *b1_mean* as well as an increase in anisotropy for the b1 shell. This pattern of change is better aligned with a positive change in *s*_*ex*_ than in any other parameter.

Figure 11b shows the estimated probability of change 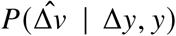 for each parameter of the standard model for an axial slice of the brain in voxels that included more than 10 WMH samples(subjects). These probabilities are normalized to sum up to 1 for each voxel. The colors indicate the probability that a change in the corresponding parameter can explain the observed changes in WMHs.

Figure 12a shows the best explaining model of change in each voxel in a few axial slices of the brain. To check for the reproducibility of the results, we have divided subjects in two batches of equal size (1500 each) and repeated the whole pipeline. The inferred changes were highly similar in the two batches with average error of 0.4% in the estimated probability of change.

**Figure 12.**
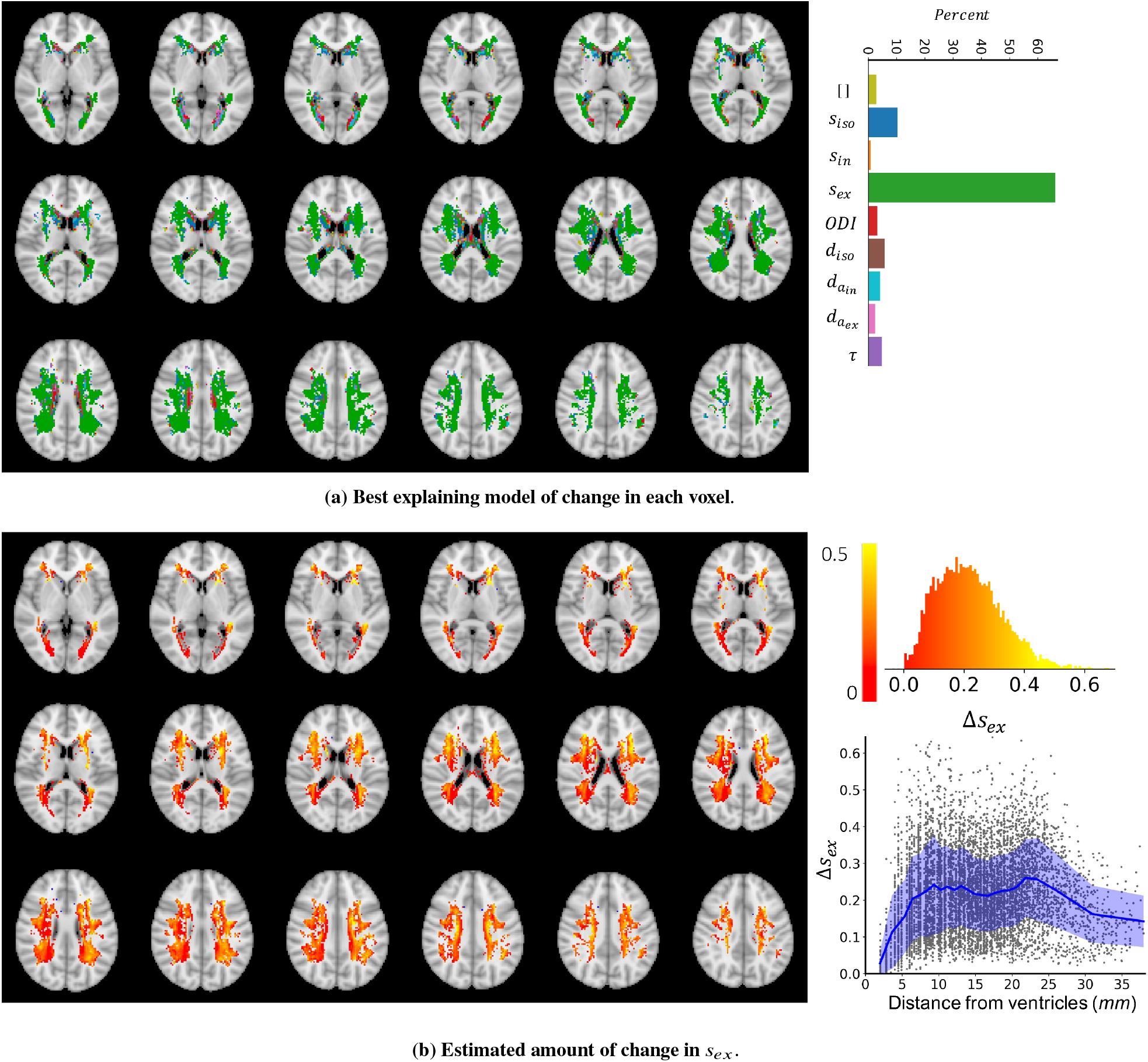
**a)** The colors indicate which model of change could best explain, i.e. had the highest posterior probability given the observed change in the summary measurements between WMH and NAWM. In the majority of voxels (65%) a change in *s*_*ex*_ explained the data better than any other model. However, in the regions very close to the ventricles there is no major winning model. This can be either because of high between subject variability or a different type of change that is not captured by the trained models of change. **b)** The maps show the estimated amount of change in *s*_*ex*_ in voxels where *s*_*ex*_ was the best model using a maximum a posteriori estimation 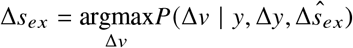. At most of the voxels the estimated amount of change is positive, meaning that an increase in *s*_*ex*_ can explain the change in the summary measurements observed in the WMH voxels. The top right panel shows the distribution of estimated amount of change at the voxels where change in *s*_*ex*_ was the best model. Most of the estimated changes are between 0 and 0.4. The bottom right panel shows the amount of change vs the distance (in millimeters) from the ventricles.

In more than 65% of the voxels, that are mostly in deep white matter, the best model is a change in *s*_*ex*_. However, in voxels adjacent to the ventricles, all other models compete and there is not a dominantly winning model. This might be due to a true difference in microstructure in these periventricular voxels, or may be caused by high variability across subjects due to CSF partial volume effects.

Figure 12b shows the estimated amount of change in *s*_*ex*_ in voxels where this was the most probable parameter. In most of the voxels an increase in *s*_*ex*_ between 0 and 0.4 explains the observed change in WMH. The bottom right panel shows that the amount of change increases with distance from the ventricles, whereas in deep white matter the average amount of change remains relatively constant.

## DISCUSSION

We presented a Bayesian framework to directly infer changes in parameters of a biophysical model from observed changes in a set of measurements. We applied the method to microstructural modelling of diffusion MRI, where biophysical models usually require many free parameters and are often degenerate.

### Comparison with model inversion

The traditional approach to overcome these degeneracies is to constrain some of the parameters to biologically plausible values so that other parameters can be estimated using a conventional measurement (e.g., fixing the diffusivities in NODDI, (Zhang et al. 2012)). Such assumptions reduce the full model parameter space to a restricted subspace, where the model is invertible. This direct inversion approach has the advantage that it gives parameter estimates and that it can model any parameter change in this restricted subspace. However, violation of these assumptions can significantly bias the parameter estimates.

Our proposed approach allows the initial set of parameters to lie anywhere within the full model parameter space (restricted only by broad user-defined priors); and any of these parameters might change. This extra flexibility comes at the price that the parameter changes are assumed to lie along 1D lines in parameter space defined by the user-provided patterns of change 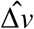. For each of these hypothesized 1D change models, we estimate the posterior probability of such a change as well as the most likely amount given the baseline data and the change in it.

To compare this assumption with that made by direct inversion, let us consider a biophysical model with 8 free parameters. Let us further assume that, due to the limited degrees of freedom in our model, we can only fit 3 out of these 8 parameters. In this case direct inversion would require assuming that the microstructural change is limited to a subset of three parameters, i.e., a 3-dimensional subspace of the full 8-dimensional parameter space. In contrast, BENCH assumes by default that the change is caused by one out of the 8 parameters, which corresponds to the microstructural change lying in one of 8 one-dimensional lines in parameter space. This suggests that if one has prior knowledge of which microstructural parameters are likely to change, it might make sense to use direct inversion with those parameters as free parameters. BENCH would have the advantage in a more exploratory approach, where any of the underlying parameters might have changed. However note that this comparison between approaches is complicated by the fact that using model inversion requires setting a subset of the parameters to some fixed value, which might cause a bias in the free parameters if inaccurately fixed (Jelescu et al. 2016; Novikov et al. 2019b). It is important to note that the user-defined prior distributions for parameters do not directly imply a prior value for the parameters. These priors are used to train the regression models and are required to be wide enough to capture all possible underlying parameter settings. Nevertheless, using broader priors only requires more complex machine learning models that can capture the variation in the relation between the measurements and their derivatives.

In the proposed approach we train the models with simulated data once (without requiring any real data) and use the trained models to estimate the desired probabilities for any real data with the same acquisition protocol. This precomputation saves one from having to integrate over all possible initial parameters when inferring the parameter change in each voxel. Therefore, the inference on real data which only consist of a few 1d integrations for each voxel, runs much faster than the non-linear optimizations in alternative inversion approaches.

The results from simulations suggest that we are able to identify changes in signal fraction accurately for the given brain-like measurement. However, there is a considerable confusion in the diffusivities, meaning that the change in these parameters is not distinguishable from one another. In simulations, we have only accounted for measurement noise, but in real data, particularly in cross-sectional studies, between-subject variability also contributes to noise. Hence, the reported performances and sensitivity to changes in parameters in the simulations section are more reliable when the between-subject variability is less important, for instance, in longitudinal studies. These accuracy values depend on the baseline measurements, underlying parameters, and the nature of how each parameter affects the measurements. Nevertheless, an important point is even in the case of full confusion in diffusivities, the results from the proposed approach is more reliable compared to the model inversion with fixed parameters. That is because a wrong prior for the fixed parameters can bias the estimates for other parameters, while in the proposed approach we avoid such assumptions. For example, in NODDI any changes in the B0 signal are usually ignored (as a result of the sum constraint on the signal fractions), but in our approach we allow changes in the b0 signal to inform which microstructural parameter might have changed.

In this paper we showed that setting a different value for diffusivities in NODDI can result in contradictory inference about changed parameters in white matter hyper intensities. The only consistent change was a decrease in the ratio of intra and extra axonal signal fractions which is in line with the results of BENCH (an increase in *s*_*ex*_ with no change in *s*_*in*_). This analysis thus illustrates one of the main benefits of using BENCH: the results do not depend on some prespecified value of a parameter as we integrate over all possible values for the parameters rather than fixing them. Another advantage is that BENCH can provide a more specific explanation for the change, e.g. in this case as opposed to NODDI that only identifies a change in the ratio of the signal fractions, BENCH can specifically tell if it is a change in the extra axonal signal fraction.

The fact that the approach doesn’t require the models to be invertible makes it applicable to studying changes in over-parameterised models or models without closed form analytical solution, e.g. simulation-based models. Such simulation-based models provide the opportunity to explore more complex and realistic models of diffusion in a tissue. There is no limitation in the number of parameters as long as they affect the observed data in some way. If several parameters cause the data to change in the same (or very similar way), this approach will give a list of possible parameters underlying the observed change with a probability associated with each. The resulting probability estimates can be used to eliminate unlikely change scenarios.

We utilized the trained models of change for the parameters of the “standard” model for diffusion to investigate which microstructural changes can explain white matter hyperintensities. The results suggest that the change can be associated with an increase in the extracellular signal. This is in line with other findings using more complex diffusion encodings (Lampinen et al. 2019), who found an increase in the extracellular T2, which would lead to an increase in the extracellular signal contribution. Comparing with the inversion approach, here we did not assume diffusivities are fixed in various brain regions, but we assumed only one of the parameters has changed as a result of white matter hyperintensity. However, it is possible that simultaneous changes in multiple parameters can better explain the change in the data, which could be tested in the same framework with the extended models of change. For example, a model with combination of the parameters might be able to explain a positive change in *b0_mean* and a negative change in *b2_mean* as it was observed in some voxels. Furthermore, we are limited to detect any changes within the constraints of the “standard” model. Hence, any changes in the signal in the white matter hyperintensities due to phenomena not within the “standard model” (e.g., exchange or non-Gaussian diffusion) would be misinterpreted as changes in the “standard” model parameters.

### Summary measures

The choice of summary measurements to train change models is arbitrary, but this choice can affect the performance of the model. It is essential that the summary measurements are able to capture enough information from the data such that they are sensitive to changes in the parameters of interest and insensitive to other changes that are not part of the model parameters. For example, in our simulations we did not include the fibre orientation parameters as part of the free parameters, and therefore we required the summary measures to be rotationally invariant. Hence the choice of decomposing the signals in each shell into spherical harmonics to extract rotationally invariant summary measurements. Of course one can instead use other signal representations, such as measures derived from the diffusion tensor model, or the kurtosis tensor model, etc, to compute the summary measurements. We chose spherical harmonics over other choices as they are fast to calculate, and the bases are orthogonal which leads to summary measures that capture different aspects of the data.

### Future developments

While in the examples shown here these patterns of change only altered a single parameter at a time, in the current framework the pattern of change can be any vector in parameter space. In the future we plan to extend this framework to allow for parameter changes in 2D or 3D hyperplanes rather than just along 1D lines (see Appendix A for the feasibility of this extension). However, the dimensionality of these hyperplanes will always be lower than that of the restricted parameter subspace in which parameters can freely change with the direct inversion approach. Note that computing posterior probabilities in a full Bayesian framework allows for comparison between models of change with different complexities without the need for arbitrary regularisation.

In addition, the model of change can be extended to study continuous changes (e.g. ageing), as opposed to discrete group differences as shown in this work. To do so, one first needs to compute the gradient of change in the measurements with respect to the independent variable, e.g. time, using a regression model. Then one can use the chain rule to relate the rate of change in the measurements to the rate of change in the parameters. Such an approach makes modelling continuous change a straightforward extension of this framework.

Although here we mostly show how our method can be applied to detect changes in parameters given the data, our framework can also be used to optimize data acquisition protocols for detecting changes in particular parameters of interest. For example, in the simulations we show that it is difficult to detect a change in the free-diffusion parameter. Our framework can be used to extend the acquisition (e.g. by adding lower bvalues) and, using the output confusion matrices, establish an optimal set of b-shells to enable detection of change in free diffusion.

Finally, while we applied the framework to the specific problem of studying microstructural changes using diffusion MRI in the brain, the framework is general meaning that it can be applied in any field where biophysical models are available. For example, the same approach as described in this paper can be applied to dynamical causal models (DCM) (Friston et al. 2003) for fMRI or MEG/EEG. These are notoriously over-parameterised, but often, are applied in a context where the values of the inferred parameters is of lesser interest than the change in the parameters under different experimental conditions, and its reasonable to assume the change is sparse; the ideal scenario for BENCH.

### SOFTWARE

BENCH is an open source software implemented in python and available at https://git.fmrib.ox.ac.uk/hossein/bench.

## ACKNOWLEDGEMENTS

SJ is supported by a Wellcome Senior Fellowship (221933/Z/20/Z), MC and SJ by a Wellcome Collaborative Award (215573/Z/19/Z). The Wellcome Centre for Integrative Neuroimaging is supported by core funding from the Wellcome Trust (203139/Z/16/Z). LG is supported by the National Institute for Health Research (NIHR) Oxford Health Biomedical Research Centre (BRC). UK Biobank Resource under Application 8107 is used in this research. We are grateful to UK Biobank for making the data available, and to all the participants, who made this resource possible by donating their time. The computations were carried out using the Oxford Biomedical Research Computing (BMRC) facilities; a joint development between the Wellcome Centre for Human Genetics and the Big Data Institute that is supported by Health Data Research UK and the NIHR Oxford Biomedical Research Centre. We additionally thank Amy Howard, Paul McCarthy, Mark Woolrich, Karla Miller, Mauro Zucchelli, and Markus Nilsson for their helpful discussions.

## APPENDIX A. TOY EXAMPLE: INFERRING CHANGES IN 2D

Consider the forward model

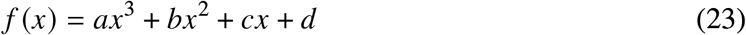

The model has 4 free parameters (*a, b, c, d*). Given 3 measurements this model is degenerate, i.e., one cannot estimate all the parameters uniquely. Now consider two instances of this model with parameters (*a*_1_, *b*_1_, *c*_1_, *d*_1_) and (*a*_2_, *b*_2_, *c*_2_, *d*_2_) with 3 measurements for each. Obviously, this system is degenerate and parameter estimation is ill posed. However, if we are only interested in comparing two model instances, we can still infer changes by assuming that the change is sparse. This is the premise of BENCH.

Now we will demonstrate that despite the model degeneracy, we can not only detect changes in a single parameter, but also infer simultaneous changes in pairs of parameters. Consider (*a*_1_ = 1, *b*_1_ = 1, *c*_1_ = 1, *d*_1_ = 1) and (*a*_2_ = 1.2, *b*_2_ = 0.8, *c*_2_ = 1, *d*_2_ = 1), i.e., Δ*a* = +0.2, Δ*b* = −0.2, Δ*c* = Δ*d* = 0.

When using Monte Carlo simulations to infer parameters for each model given three independent measurements, the posterior distribution is clearly degenerate as shown in Figure 13a. In this figure, the blue (resp. red) distribution shows the parameter estimates for (a, b) for the first (resp. second) data set. The intensity of each point encodes the log posterior probability for the estimated parameters. The stars show the true parameter values. The plot demonstrates that parameter estimates are highly correlated (i.e. the model is degenerate).

**Figure 13.**
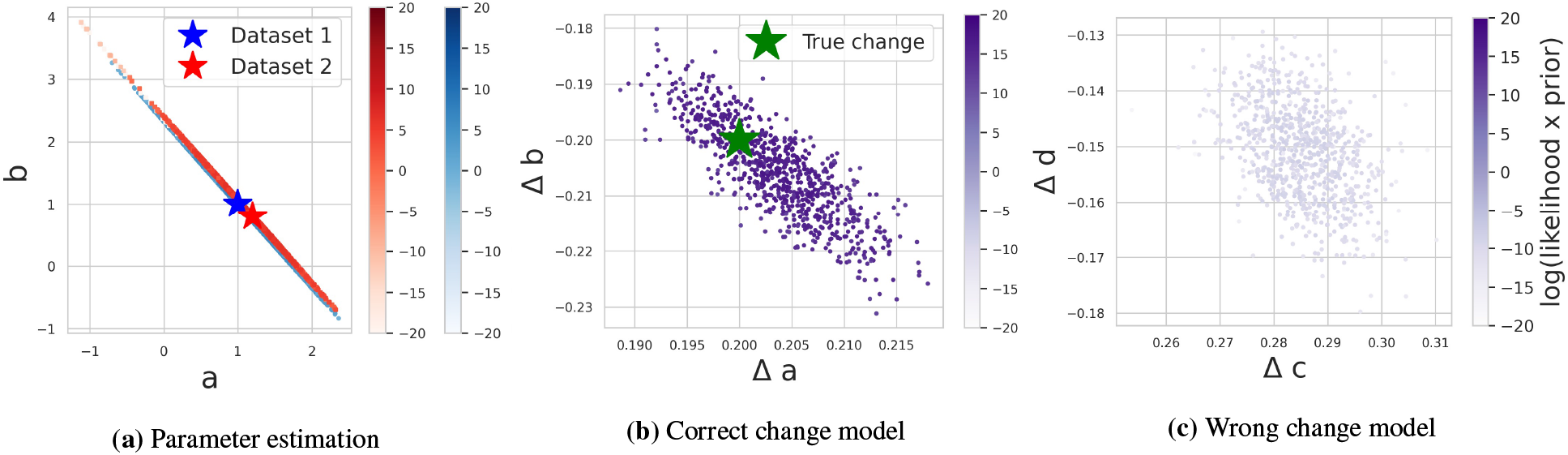
a) Parameter estimation. Each set of dots shows parameter estimates for one instance of the model using MCMC and intensities represent the log posterior probability. The parameter estimates for each data set are highly correlated and all of the points on the lines explain the data equally well, i.e. the models are degenerate and it is not possible to directly compare the parameter estimates. B) Inferred change with the correct model. We ran MCMC with the assumption that change has a particular shape (only a and b changed). The estimated values for Δ*a* and Δ*b* are centred around the correct change (green star) and the unnormalized posterior probabilities are comparatively high. C) Inferred change using a wrong model. We run a similar MCMC but this time assuming *c* and *d* can change. In this case the estimated posterior probabilities are much smaller compared to the previous change model, i.e. this model of change cannot explain the change in the measurements as well as the model in (b).

In contrast, figure 13b shows Monte Carlo samples for Δ*a* and Δ*b* for the change model. The plot demonstrates that the estimated parameter changes are distributed around the true change value and each sample has a comparatively high posterior probability value. It is therefore possible to infer the true, 2-dimensional change.

We also considered an alternative change model where a and b are fixed and c and d can change. The estimated samples for Δ*c* and Δ*d* are shown in Figure 13c. In this case the estimated samples have much lower posterior probabilities (lower intensities) than the a,b change model. Thus, we can use the change model to assess not only the amount of change in 2D, but also which pair of parameters best explains these changes. The changes are still sparse, but not necessarily 1-dimensional.

In BENCH we integrate the approximations of this unnormalized posterior probabilities to compute the the desired probabilities for each model of change in Eq. 1. Hence, it this example BENCH (once extended to allow multi-dimensional changes) would correctly infer that it was the parameters a and b that changed, and not the parameters c and d.

## APPENDIX B. ESTIMATING QUALITY OF FIT

The estimated probability in Eq.1 tells how well each model explains the observed change compared to all other defined change models, but it doesn’t necessarily reflect to what extent the observed and predicted change are matched. In other words, a model with a poor quality of fit to the data can get a high probability value because its prediction is the closest to data compared to all other models. Also, it is possible that more than one change model predict the data accurately and hence all get low probabilities in Eq.1.

To estimate how well a change model can explain the change in data one can look at the chi-squared distance between the predictions of the change model and the measured change:

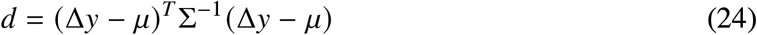

In the above expression, Δ*y* is the observed change in the data, and *µ* and Σ are the mean and covariance of change in the measurements predicted by the best model. This statistic follows a chi-squared distribution and a higher *d* means more discrepancy between the observed change and the predicted change.

Figure 14 shows the distribution of *d* for the case of one parameter change that is explained by the correct model (blue) and the case of two parameter change that is mistakenly identified as a single parameter change (orange). Accordingly, our recommendation when the discrepancy is high is to consider revising the change models, as the winning model is poorly explaining the observed change. For example, one can define biologically feasible linear combinations of the parameters as change directions.

**Figure 14.**
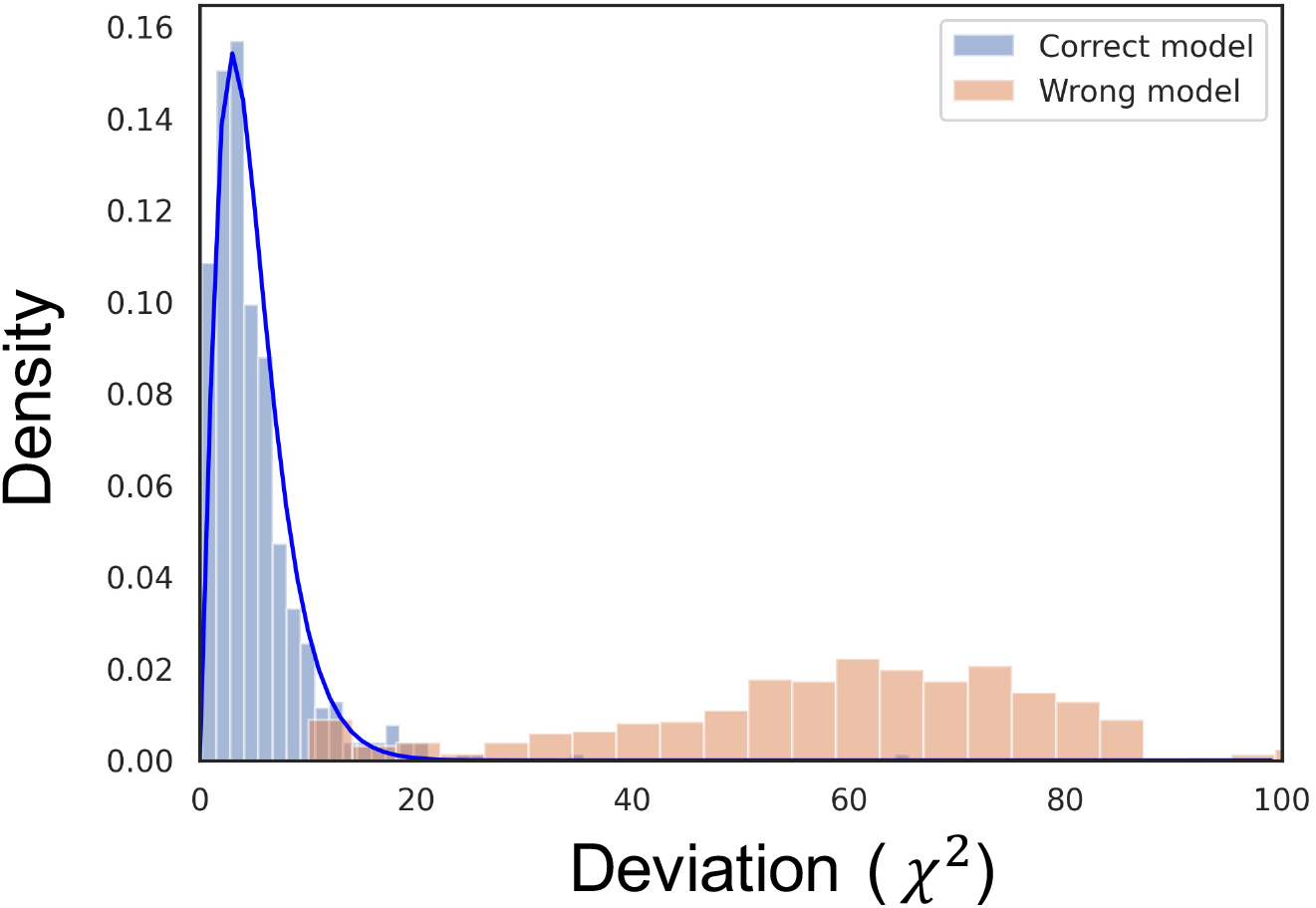
Distribution of distance for the correct change model (blue) and a wrong model(orange). Given a baseline measurement (*y*) and a change (Δ*y*), we estimate the most likely change in the parameters as well as the most likely amount of change in that direction using our trained change models. These estimates can then be used to predict the distribution of expected change in the measurements. Using the discrepancy between this prediction and the actual observed change, we can determine the quality of the change model in explaining the data. The histograms are showing the Mahalanobis distance (i.e., the offset normalised by the covariance matrix as defined in 24) between the actual and the predicted change in the measurement when the correct change model is used (blue) and when a wrong change model is used (orange) for several instances of simulated data. The blue curve shows the pdf of χ^2^ distribution with *df* = the number of measurements.

